# Brain-wide measurement of synaptic protein turnover reveals localized plasticity during learning

**DOI:** 10.1101/2022.11.12.516226

**Authors:** Boaz Mohar, Gabriela Michel, Yi-Zhi Wang, Veronica Hernandez, Jonathan B. Grimm, Jin-Yong Park, Ronak Patel, Morgan Clarke, Timothy A. Brown, Cornelius Bergmann, Kamil K. Gebis, Anika P. Wilen, Bian Liu, Ricard Johnson, Austin Graves, Tatjana Tchumatchenko, Jeffrey N. Savas, Eugenio F. Fornasiero, Richard L. Huganir, Paul W. Tillberg, Luke D. Lavis, Karel Svoboda, Nelson Spruston

**Author notes:** Co-Senior Authors (Karel Svoboda, Nelson Spruston).

## Abstract

Synaptic plasticity underlies learning and memory by altering neuronal connections in response to experiences^1^. However, the loci of learning-induced synaptic plasticity, and the degree to which plasticity is localized or distributed, remain largely unknown^2^. We developed a method (DELTA) for mapping brain-wide changes in synaptic protein turnover with single-synapse resolution, based on Janelia Fluor dyes and HaloTag knock-in mice. During associative learning, the turnover of the ionotropic glutamate receptor GluA2, an indicator of synaptic plasticity, was enhanced in several brain regions, most markedly in the hippocampal area CA1. More broadly distributed increases in turnover of synaptic proteins were observed in response to environmental enrichment. In CA1, GluA2 stability was regulated in an input specific manner, with more turnover in layers containing input from CA3 compared to entorhinal cortex. DELTA will facilitate exploration of the molecular and circuit basis of learning and memory and other forms of adaptive and maladaptive plasticity at scales ranging from single synapses to the entire brain.

## Main

Cellular functions are tuned by protein synthesis and degradation, which results in dynamic protein turnover^3–5^. In the brain, protein lifetimes range from tens of minutes for immediate early gene proteins^6^ to months for a subset of synaptic structural proteins and many others^7–9^. Turnover rates vary by protein, cell type, and brain region^10–12^. Protein dynamics also depend on environmental and behavioral conditions^12,13^ that are necessary for animals to learn cognitive and motor tasks^14–16^, and are dysregulated in neurodegenerative diseases^17^. Learning is supported by long-term changes in synaptic strength that require both protein synthesis^18,19^ and degradation^20–22^. Since behavior-related plasticity occurs in a coordinated manner across multiple brain regions^2^ but may be limited to specific synapses in individual neurons^1,23,24^, measurements of protein turnover are needed at multiple spatial scales, ranging from brain wide to subcellular.

Metabolic incorporation of stable isotope labeled amino acids followed by mass spectrometry (MS) allows the measurement of turnover for many proteins in parallel^25^, but its spatial resolution is limited to the level of brain regions (*e.g.*, cortex *vs.* cerebellum) or homogenate fractions (*e.g.*, cytoplasm *vs.* synaptosomes). Recently, a suite of new tools^26–29^ has enabled proteins of interest to be labeled with fluorescent markers and imaged with high contrast and resolution, including a pulse-only^28^ method using the HaloTag^30^ (HT) and fluorescent ligands. For an in-depth comparison of these methods, see Supplemental Text.

Here, we identified a panel of HaloTag ligand (HTL) Janelia Fluor (JF) dyes^31–34^ that are optimized for brain-wide pulse-chase labeling of synaptic proteins in HT knock-in mice. We used these JF-HTL dyes to develop DELTA (**D**ye **E**stimation of the **L**ifetime of pro**T**eins in the br**A**in), a method that measures protein lifetimes (τ) *in vivo* using pulse-chase experiments with spectrally separable fluorescent ligands. DELTA enables precise measurements of protein lifetime in the whole brain and other tissues *in vivo,* down to synaptic resolution. We applied DELTA to three knock-in mouse lines expressing the HT protein, including a newly generated glutamate receptor subunit (GluA2) HT knock-in mouse, a predominant AMPA-type glutamate receptor subunit^35^, which allowed us to map learning-related synaptic plasticity on a brain-wide scale.

### Modeling and measuring protein turnover *in vivo* with high spatial and temporal resolution

We first modeled the HT dye-ligand labeling process to identify conditions under which a pulse-chase method would enable precise protein turnover measurements *in vivo* of a HT-fused protein (**Fig. 1a**). We considered three populations of HT fusion proteins: (1) *Unlabeled* protein– HT; (2) *Pulse*, the population labeled with the first dye-ligand infusion; and (3) *Chase*, the newly synthesized protein-HT population labeled with a second infusion using a spectrally distinct dye-ligand. Assuming an exponential decay of these protein populations and stable overall protein concentrations, we can calculate their mean lifetime τ = Δ*t*/log(1/*Fraction Pulse*), where Δ*t* is the time interval between pulse and chase dye administration and *Fraction Pulse* is the proportion of protein-HT fusion labeled by the pulse dye-ligand (*Fraction Pulse* = *Pulse* / [*Pulse* + *Chase*]). As slow dye clearance effects turnover measurements (gray shaded region in **Fig. 1a**; **Supplementary text**), we measured dye-ligand clearing using *in vivo* imaging and determined an effective time constant of approximately 82 min (**Extended Data Fig. 1**).

**Figure 1.**
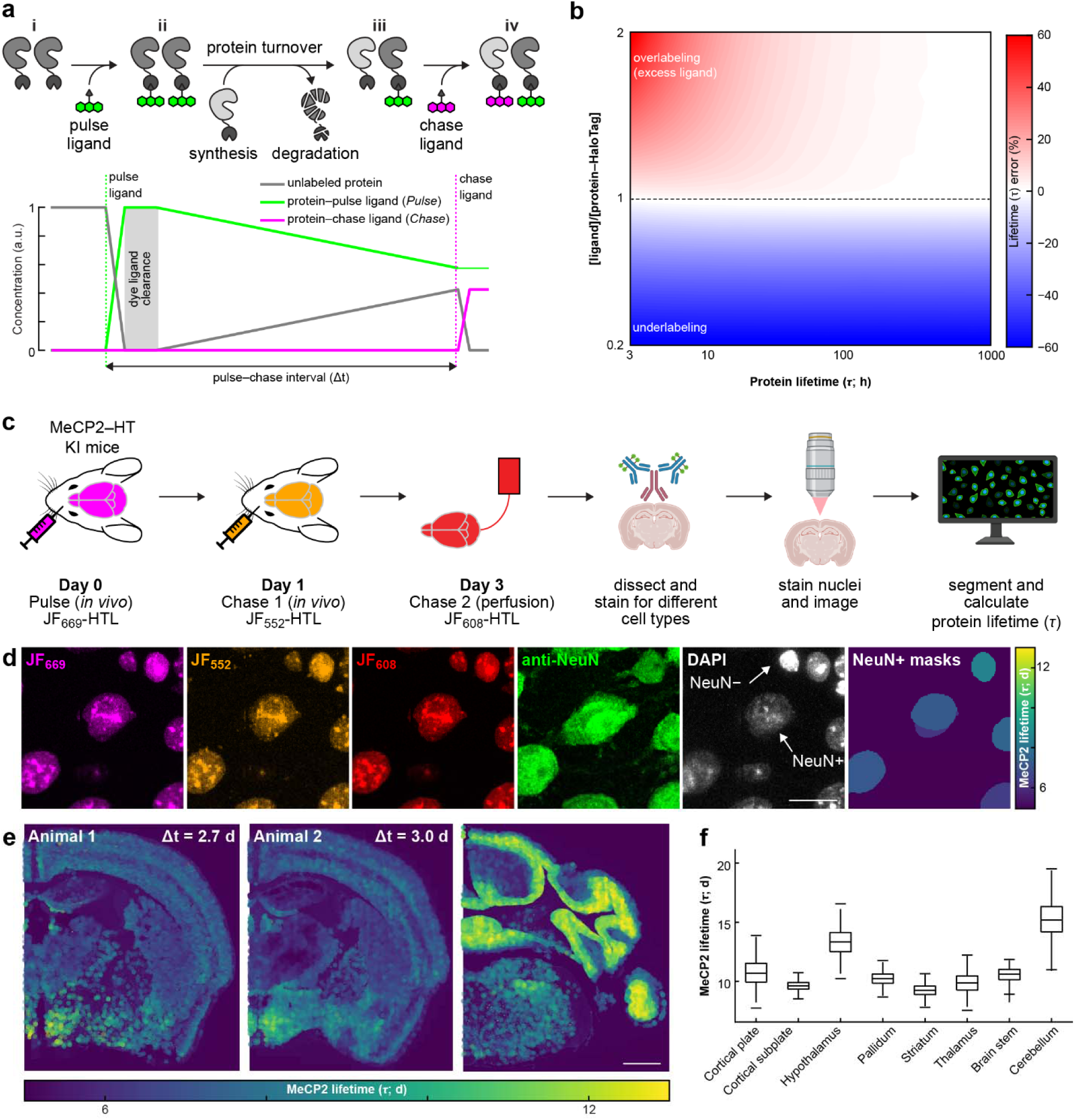
Measurement of protein turnover *in vivo*. **a,** Schematics showing how DELTA can measure protein lifetime by sequential HTL dye capture using a HT modified protein. (i) Before the injection of the first dye ligand (pulse dye), all proteins are unlabeled (gray line). (ii) After injection of the pulse dye (dashed green line), all proteins are labeled with the pulse dye (solid green line - *Pulse*). (iii) During the pulse-chase interval, some proteins degrade, and others are synthesized but are unlabeled. (iv) Injection of a spectrally separate dye (dashed magenta line) binds the newly synthesized protein (solid magenta line - *Chase*). The gray shaded area indicates where excess dye delays the onset of turnover measurement, leading to a *Pulse* overestimation error (see also Supplementary Text). **b**, The estimated lifetime error (color) as a function of dye-protein ratio (y-axis) and true protein lifetime (x-axis). Undersaturation (<1 ligand/protein ratio) causes worse errors than dye excess and longer-lived proteins are estimated more accurately than short-lived ones. **c-d,** Turnover measurement of the nuclear protein MeCP2-HT in a knock-in mouse model. (c) Experimental design: Three HTL dyes were used to measure multiple protein-turnover intervals. Additionally, after perfusion and dissection, coronal sections were labeled with DAPI and fluorescent antibodies to identify different cell types. (d) Example field-of-view showing the JF dyes (left 3 panels) with NeuN and DAPI for identification of neuronal nuclei. After segmentation of NeuN positive nuclei, segmented nuclei were colored by lifetime using the sum of the two *in vivo* injections as the *Pulse* (*Fraction Pulse* = [JF_669_+JF_552_]/[JF_669_+JF_552_+JF_608_]). **e,** Example coronal sections from two animals. Left and middle, illustration of the consistency of the lifetime estimates for aligned AP sections; Right, longer lifetime in the cerebellum (compared to middle and left panels). **f,** MeCP2-HT neuronal nuclei lifetime (bootstrap of means from 5 animals and 3 intervals where line is median, box is 25-75^th^ percentile and whiskers are 0.5-99.5^th^ percentile) across CCF-aligned brain regions.

Using modeling and *in vivo* dye-ligand clearance measurements, we investigated the effects of the three experimentally controlled variables on estimates of lifetime using DELTA: (1) the selection of a target protein, which determines mean lifetime; (2) the amount of pulse dye injected, which determines ligand to target ratio; and (3) the pulse-chase interval (Δ*t*; **Fig. 1b**; **Extended Data Fig. 2; Supplementary Text**). We found that underlabeling (less dye-ligand than target protein) produced large errors, regardless of lifetime. However, dye-ligand excess produced only small errors over a large range of lifetime values, except for proteins where lifetime is on the order of or shorter than dye-ligand clearance (**Fig. 1b**). As most proteins in the brain have lifetimes far longer than dye clearance times (lifetimes days to weeks; effective clearance time constant ∼82 min)^3,12^, dye-ligand excess is the preferred regime. Additionally, dye-ligand excess would not distort relative measurements of protein lifetime across cell types, brain regions, or individual animals. Compared to single-dye methods (pulse-only)^28^, the normalization provided by the chase ligand in DELTA increased signal-to-noise ratio (SNR) to measure turnover (**Extended Data Fig 2g, h; Supplementary Text**). Changes in total protein levels can cause errors in lifetime estimation using the approach outlined above. However, such changes can be detected (estimated by summing the *Pulse* and *Chase*) and are easily corrected (**Extended Data Fig 2i**).

The DELTA protocol requires fluorescent HTL dyes that are spectrally distinct and efficiently bind HT fusion proteins *in vivo*. We screened for suitable JF HTLs^31–34^ in mouse brains and identified JF_669_-HTL and JF_552_-HTL as particularly bioavailable, with both HTL dyes having strong and uniform labeling across the brain (**Extended Data Fig. 3, 4; Supplementary Text)**. We subsequently tested a more photostable dye, JFX_673_-HTL^36^, which retained the high bioavailability of its parent JF_669_-HTL^32^. Finally, we investigated the optimal *in vivo* dye-ligand formulation and delivery protocols (**Methods**; **Extended Data Fig. 5,6; Supplementary Text**).

We first used DELTA to measure the lifetime of the nuclear protein methyl-CpG binding protein 2 (MeCP2)^37–39^ using MeCP2–HT knock-in mice^40^. As MeCP2 is relatively abundant (top 10% in the brain^41^), achieving dye excess would signify DELTA’s broad applicability (**Extended Data Fig. 7**). Previous MeCP2 lifetime measurements using MS also provide a point of comparison for our method^12^. We compiled multiple estimates of protein lifetime from the same animal by sequentially injecting the highly bioavailable dye-ligands JF_669_-HTL ligand and JF_552_-HTL, followed by perfusion of a third dye-ligand: JF_608_-HTL. After perfusion, the brain was sectioned and stained with DAPI to identify all nuclei, and Immunofluorescence (IF) was used to classify cell types (**Fig. 1c; Extended Data Fig. 8d**). We imaged all three dyes at subcellular resolution and calculated *Fraction Pulse* for all neuronal nuclei across the brain of five MeCP2–HT mice. We converted the measurements to average protein lifetime (τ; **Fig. 1d,e**). The lifetime of neuronal MeCP2 differed across brain regions (medians ranged from 9.2 to 15.2 days; **Fig. 1f, Extended Data Fig. 8a–c**). The longest MeCP2 lifetime was in the cerebellum (**Fig. 1e**, right panel), in agreement with previous measurements using MS^12^. Also consistent with previous studies, we found that MeCP2 was expressed at higher levels in neuronal cells than in glia (**Extended Data Fig. 8d–f**)^42^. These results highlight the ability of our pulse–chase paradigm for lifetime measurements of abundant proteins at brain-wide scales.

### PSD95-HaloTag turnover is accelerated by environmental enrichment

PSD95 is an abundant scaffold protein in the postsynaptic density of excitatory synapses^43^ and anchors glutamate receptors in synapses. Synaptic PSD95 content is known to be regulated by protein synthesis and degradation^44,45^. We measured brain-wide PSD95 lifetimes and the effects of behavioral manipulations in synapses using knock-in mice expressing PSD95 fused with HaloTag (PSD95–HT^46^; **Extended Data Fig. 9**). Mice housed in standard cages (*n* = 4) and mice placed in an enriched environment^47,48^ (EE; *n* = 4) were imaged with a pulse–chase interval of 14 days, using JFX_673_-HTL as the pulse injected *in vivo* and JF_552_-HTL as the chase in perfusion (**Fig. 2a**). This order of dye-ligands ensures saturation of all expressed proteins due to the high bioavailability of JFX_673_-HTL, which is similar to its parent ligand, JF_669_-HTL (**Extended Data Fig. 3j, 6d)**. Following imaging (**Fig. 2b,c**), sections were registered to the Allen Brain Reference Atlas (CCFv3)^49^, and the lifetime of PSD95-HT was calculated (**Fig. 2d,e**). PSD95-HT lifetimes in control animals varied 50% across brain regions and cortical layers (10.9 – 17.8 days; **Fig. 2f,g**). The brain-wide average lifetime of PSD95 was more than two days shorter (∼21%) in mice housed for two weeks in EE compared to control (**Fig. 2h**). The neocortex showed the largest reduction in PSD95 lifetime with EE (∼30%; **Fig. 2i**). Effects of EE on lifetime were similar across cortical layers but not hippocampal subfields (**Fig. 2j**). We detected small increases in total protein in some brain regions after EE. DELTA allowed us to correct for this increase in our estimate of protein lifetime (**Methods**; **Extended Data Fig. 2i**).

**Figure 2.**
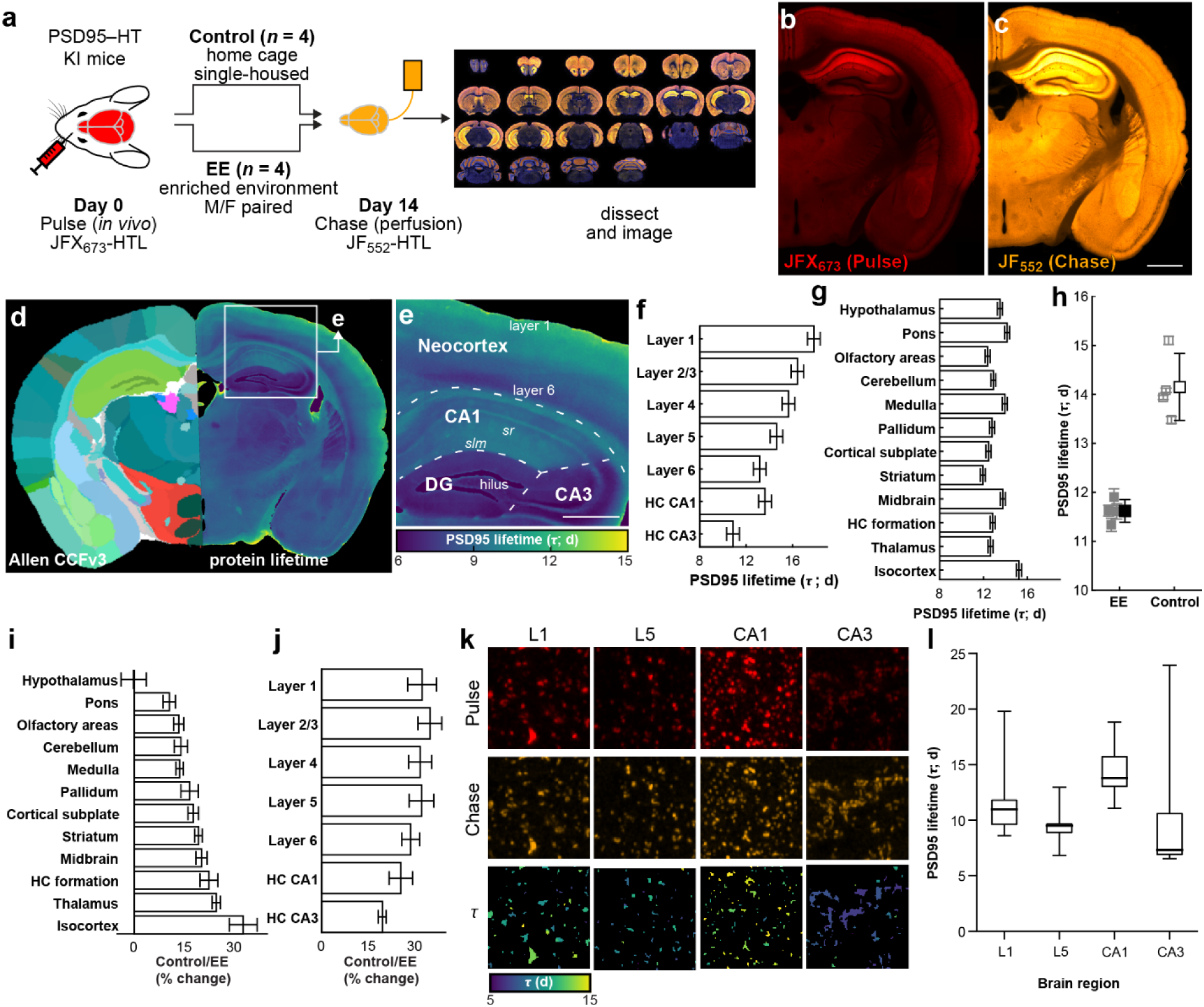
PSD95 turnover depends on experience. **a,** Experimental design for turnover measurement of PSD95-HT in a knock-in mouse. See text for details. **b-e,** Example coronal section showing the *Pulse* (b), *Chase* (c), and calculated lifetime aligned to the Allen CCFv3 (d). **e,** zoom-in of the image from panel (d). Note the lifetime gradient that separates the CA1 *stratum moleculare (sr), stratum lacunosum-moleculare* (*slm*; long lifetime) from dentate gyrus (DG) and hilus (all short lifetime). **f**, Average lifetimes of control animals (*n* = 4) cortical layers and subfields of the hippocampus (HC) are significantly different (Mixed-effects linear model with layers as fixed effects [10-16.5 days means with ∼0.5 SE all *p* < 0.0001] and animal ID as a random effect with mean of 1.42 days and mean residual error of 1.39 days). **g**, Average lifetime for twelve large brain regions is also significantly different in Control mice (*n* = 4; Mixed-effects linear model with brain regions as fixed effects [11.9-15.2 days means with ∼0.23 SE all *p* < 0.0001] and animal ID as a random effect with mean of 0.46 days and mean residual error of 1.8 days). **h,** Average lifetime of four mice under EE and four under Control conditions. EE increased protein turnover and shortened the average lifetime of PSD95-HT. Individual animals in lighter colors (EE: 11.6±0.2 days *n* = 4; Control, green: 14.2±0.7 days *n* = 4; Wilcoxson rank sum test *W* = 26, *p* = 0.0286). **i,** Percent change in Control vs. EE animals for twelve different brain regions; Mixed-effects linear model for each brain region with group assignment as fixed effects and animal as a random effect. **j,** same as (i) for cortical layers and hippocampus (HC) subfields. **k,** Example images using Airyscan imaging of ExM tissue (max projection of 5 z planes 0.3 um apart) from layer 1 (top row), HC CA1 subfield (middle row), and basal dendrites of CA3 (bottom row) showing both *Pulse*, *Chase*, and lifetime. **l,** Quantification of segmented single synapse turnover (median lifetime in days [lower quartile – upper quartile]: L1 10.97 [9.62-11.79], L5 9.5 [8.89-9.66], CA1 13.78 [13.04-15.72], CA3 7.32 [6.92-10.59]).

PSD95 lifetimes measured by the pulse-chase DELTA were substantially longer than those measured using HT ligands in a pulse-only approach^28^ (e.g., for control neocortex mean [CI]: 15.2 [14.7-15.6] days for DELTA vs. 8.1 [6.1-12.0] for pulse-only), even though both methods are based on the same PSD95-HT knock-in mice (**Extended Data Fig. 9f,g)**. To obtain an independent estimate we turned to stable heavy isotope-based MS measurements of protein turnover. We fed either wild-type or homozygous PSD95-HT knock-in mice a ^13^C_6_-lysine diet for 7 or 14 days (**Extended Data Fig. 9h**; four animals per group for each pulse interval; *n* = 16 animals in total). The neocortex was then isolated and protein lifetime was measured^12^. We detected a small difference in the lifetime of PSD95 in wild-type (WT) vs. HT knock-in mice (mean [95% CI] in days: WT: 19.3 [17.1-21.6], HT knock-in: 15.9 [13.3-18.6], t-test *t*(8) = 2.2268, *p* = 0.056, *n* = 4 for each). The MS measurements showed turnover rates close to DELTA (15.2 [14.7-15.6]), but longer than estimates using the pulse-only method^28^ (**Extended Data Fig. 9i**). The MS measurements also showed that most other proteins did not change in their turnover rate in the PSD95-HT knock-in animals, validating their use in these experiments (**Extended Data Fig. 9j; Supplementary Table 3**).

To investigate whether DELTA could be used to measure PSD95-HT lifetime in individual synapses, which requires approximately four-fold higher imaging resolution than permitted by the diffraction limit^50^, we developed an expansion microscopy (ExM) protocol that enables two-fold expansion without proteolysis or strong protein denaturation, thus fully retaining HT-bound JF-HTL dyes. Imaging with Airyscan confocal microscopy^51^ provided another two-fold improvement in resolution, providing the required four-fold enhancement. Combined with our use of bright and photostable small-molecule dyes, this approach enabled measurements of turnover at individual synapses. We measured the turnover in individual synapses of neocortical layers 1 and 5, and hippocampal subfields CA1 and CA3 (**Fig. 2k,l**; **Extended Data Fig. 9c; Supplementary Movie 1**). Our analysis revealed that layer 1 exhibited slower turnover compared to layer 5 in the neocortex, and similarly, CA1 showed slower turnover than CA3 in the hippocampus. While these trends align with those observed in whole-brain imaging, it’s important to note that our single-synapse imaging focuses on a smaller, localized region within each structure. As a result, these localized measurements are not directly comparable to whole-brain imaging, which involves significant averaging across cellular compartments these large structures.

These results demonstrate that DELTA can accurately estimate synaptic protein lifetimes with brain-wide coverage, up to single-synapse resolution, and is sensitive to behavioral manipulations.

### Learning-related GluA2-HaloTag turnover is enriched in the hippocampus

We next used DELTA to identify brain regions with enhanced synaptic plasticity during learning. As ionotropic glutamate receptors, including GluA2, directly influence synaptic strength and are modified during synaptic plasticity^52–54^, we probed turnover in a novel GluA2-HT knock-in mice (**Extended Data Fig. 10a**). Homozygous GluA2-HT mice had normal GluA2 protein levels (**Extended Data Fig. 10b**) and showed high correlation between the HTL-dye signal and immunostaining of GluA2 (**Extended Data Fig. 10c,d**). Recordings from CA1 pyramidal neurons in acute hippocampal brain slices showed normal synaptic activity (**Extended Data Fig. 10e,f)** and long-term potentiation (**Extended Data Fig. 10g,h).**

To study the turnover of synaptic proteins associated with learning, head-restrained, water-restricted mice were trained in a virtual-reality arena. The mice were conditioned to lick for water rewards as visual cues appeared at pseudo-random locations within an infinitely extending corridor (**Methods**, **Fig. 3a**). We first trained mice to gather rewards at locations marked with one of two distinct visual cues (stripes and circles; reward probability 50% for both cues). After mice (*n* = 7) reliably licked in the presence of visual cues but not otherwise, they were injected with the pulse dye (JFX_673_-HTL). To investigate new learning, a subset of mice was switched to a task variant in which only the stripes visual cue was rewarding while the other cue was not (New Rule; *n* = 3; stripes reward probability 100%). Mice learned to lick only in the presence of the stripes visual cue, as indicated by increased licking efficiency within three days (Efficiency = # rewards / # cues licked; Example mouse in **Fig. 3b**). The other mice (Baseline; *n* = 4) remained on the initial task and maintained licking efficiency around 50%. At the end of three days with one behavioral session per day, there was a clear difference in licking efficiency between the groups, and all mice were perfused with the chase dye (JF_552_-HTL; **Fig. 3c**).

**Figure 3.**
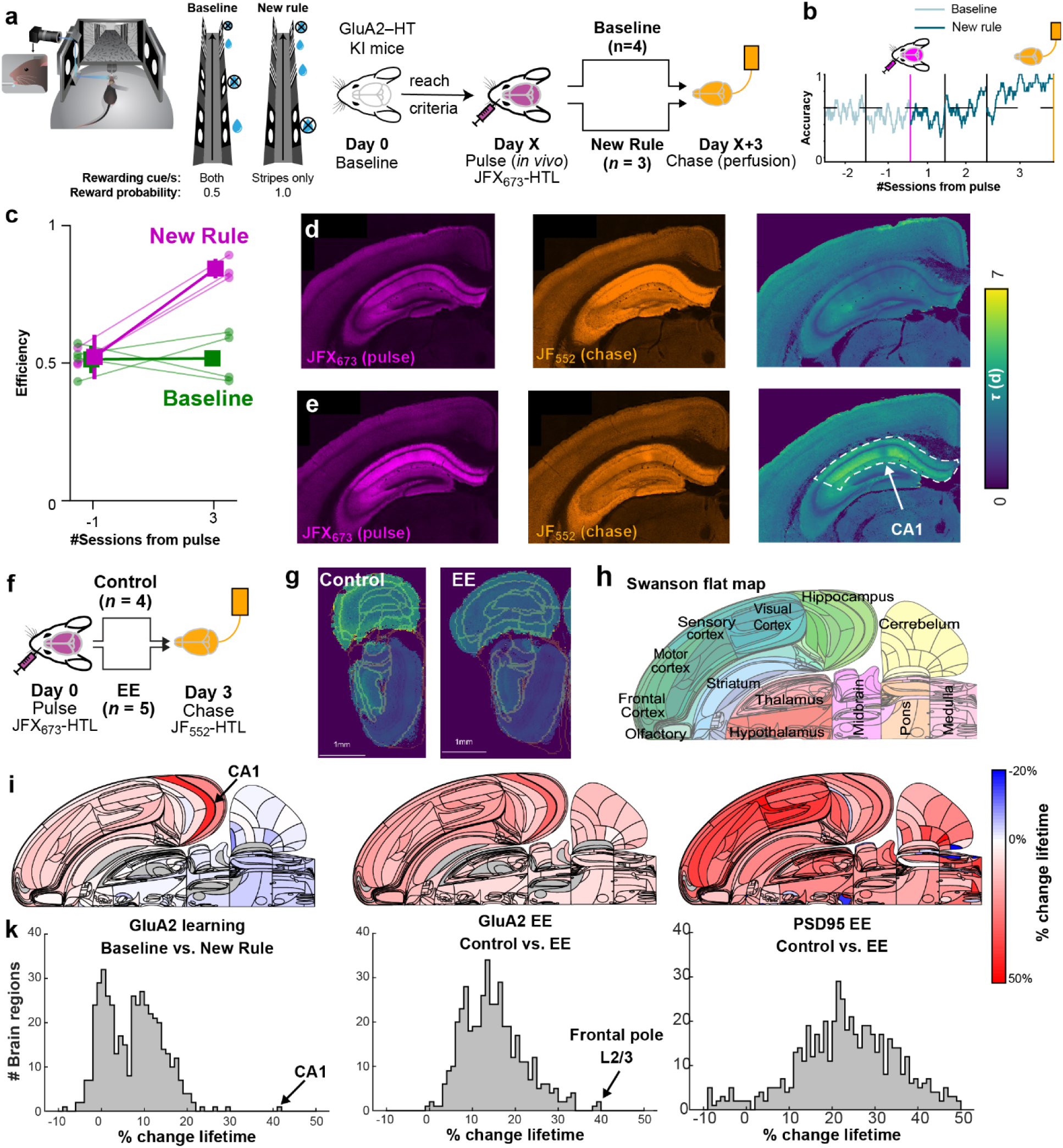
Learning-induced GluA2 turnover is more localized than following environmental enrichment. **a,** Experimental design for turnover measurement of GluA2-HT knock-in mouse during learning. See text for details. **b**, Example of efficiency (# rewards / # cues licked) across 5 sessions (1 session / day) for a mouse in the New Rule condition. For the first 2 sessions both cues are rewarding at 50% probability (light gray line) and the mouse **does not** achieve consistently high efficiency under these conditions. Turnover measurements were done by injecting pulse dye (JFx_673_-HTL) after the second session and perfusing with the chase dye (JF_552_-HTL) after 3 sessions (1 session a day). After pulse dye injection the mice were switched to the New Rule condition in which only the stripes visual cue is rewarding at 100% probability (dark gray line). By the end of the third session after the switch the mouse learn to lick only in the stripes cue and avoid licking in the circles as evident by near-perfect efficiency. **c,** Efficiency of all seven animals comparing the session before pulse dye injection (-1) to efficiency 3 session after pulse dye injection. Baseline animals (green) did not change in efficiency while New Rule animals (magenta) improved. **d**, Example coronal section from the animal in (b). *Pulse* (left panel) and *Chase* (middle panel) were imaged post-hoc across the brain and lifetime of GluA2-HT (right panel) was calculated. Notice low lifetime in CA1 region of the HC. **e**, Same as (d) for an animal that stayed in the *Baseline* condition. Notice the higher lifetime in CA1 region of the HC. **f**, Experimental design for turnover measurement of GluA2-HT modulated by EE. **g**, Example coronal sections from Control mice (left panel) and after EE (right panel). Notice the lower lifetime of GluA2-HaloTag in frontal cortical regions. **h**, Swanson flatmap^89^ representation of mouse brain regions. Different colors represent different brain regions. **i**, Left panel: % change in GluA2-HaloTag lifetime in **Baseline** vs. **New Rule** learning groups. The largest effect is in HC, specifically in CA1 (black arrow). Middle Panel: % change in GluA2-HaloTag lifetime in Control vs. EE groups. Effects are more distributed than learning. Right panel: Panel iv:. % change in PSD95-HaloTag lifetime in *Control* vs. *EE* groups. Same data as in Figure 2. **k**, Distributions of changes in turnover over the 3 groups as in (i). GluA2 learning (left panel) shows a sparser distribution with more regions with close to 0% change (*n* = 442, median = 7.5%, 59.5% of brain regions with over 5% change), while both GluA2 EE (middle panel) and PSD95 EE (right panel) have more regions with larger changes (PSD: *n* = 580, median = 23.1%, 95.9% of brain regions with over 5% change; GluA2: *n* = 442, median = 14.3, 94.8% of brain regions with over 5% change).

Brains were imaged and aligned to the Allen Brain Reference Atlas (CCFv3; **Fig. 3d-e)**. In the raw data, large changes in GluA2-HT turnover were apparent in the neocortex and hippocampus. The New-Rule group CA1 region had low *Pulse* signal (**Fig. 3d left panel** - **magenta**) and high *Chase* signal (**Fig. 3d middle panel** - **orange**) indicating fast GluA2-HT turnover (**Fig. 3d right panel - green and blue shades**). In the Baseline group, CA1 and superficial neocortical layers showed higher *Pulse* signal (**Fig. 3e left panel** - **magenta**) and lower *Chase* signal (**Fig. 3e middle panel** - **orange**) indicating slower GluA2-HT turnover (**Fig. 3e right panel - yellow shades**).

We compared the changes in GluA2-HT turnover during learning vs. EE (**Fig. 3f, Extended Data Fig. 10i-k**; Control *n* = 4; EE *n* = 5). As with PSD95-HT, EE accelerated turnover of GluA2-HT, but unlike rule learning, the largest effect of EE was in frontal cortical areas (∼40% in layer 2/3 of the frontal pole; **Fig. 3g**). We compared all three manipulations using the Swanson flat map representation (**Fig. 3h)**. The largest difference in GluA2-HT turnover in the New-Rule group was in the hippocampus (**Fig. 3i left panel**). Changes caused by EE both for GluA2-HT (**Fig. 3i middle panel**) and PSD95-HT (**Fig. 3i right panel**) were more widely distributed across brain regions. Unlike PSD95-HaloTag, total GluA2-HaloTag levels were stable across all four conditions (data not shown).

We examined the similarity of plasticity across conditions by characterizing the distributions of turnover changes. All 3 distributions were different (Kruskal-Wallis test χ^2^ = 499.4, p = 3.6e-109, Tukey-Kramer post-hoc test all medians are significantly different) with GluA2 learning show the smallest median (**Fig. 3k right panel**, *n* = 442, median = 7.5%) followed by GluA2 EE (**Fig. 3k middle panel**, *n* = 442, median =14.3%) and PSD95 EE showing the largest median (**Fig. 3k right panel**, n = 580, median = 23.1%). For EE in both PSD95 and GluA2 the distribution shows most brain regions were modulated (95.9% and 94.8% of brain regions with over 5% change), whereas for learning in GluA2 the distribution was sparser with more brain regions centered around 0% change (59.5% of brain regions with over 5% change). This data shows that the distribution of turnover varies across synaptic proteins, even in the same complex (*i.e.*, synapses), and across different behavioral manipulations.

These results indicate that DELTA can be used to identify the loci of plasticity for multiple synaptic proteins and behavioral manipulations.

### Sub-cellular regulation of synaptic protein turnover

Synaptic protein turnover was not uniform across the large spatial extent of neurons, hinting at the existence of compartment-specific plasticity. This was most apparent in the CA1 region of the hippocampus, where cell bodies reside in a narrow layer (**Fig. 4a** *stratum pyramidale*) and axons from different pathways impinge on dendrites of the same cells in different layers. CA3 axons arrive on proximal dendrites (basal, *stratum oriens, so;* apical, *stratum radiatum, sr*), whereas axons from layer 2 of entorhinal cortex innervate distal apical dendrites (*stratum lacunosum moleculare, slm*). GluA2-HT turnover exhibited a complex spatial pattern across layers (**Fig. 4b**). First, in *so* and *sr*, lifetime increased with distance from the soma (from 3.5 to 7 days in *sr*). Second, cell bodies showed very short GluA2-HT lifetimes. Third, the transition from *sr* to *slm* was marked by a steep drop in lifetime (back to 3.5 days) and a lack of distance-dependent gradient from the soma. PSD95-HT turnover showed a similar spatial pattern across CA1 layers (**Fig. 4c**) except for the fast turnover at the somatic layer. Fourth, at higher resolution, CA1 somata appeared to contain an intracellular pool of newly synthesized (< 3 days old) GluA2 but not PSD95 (**Fig. 4d,e**).

**Figure 4.**
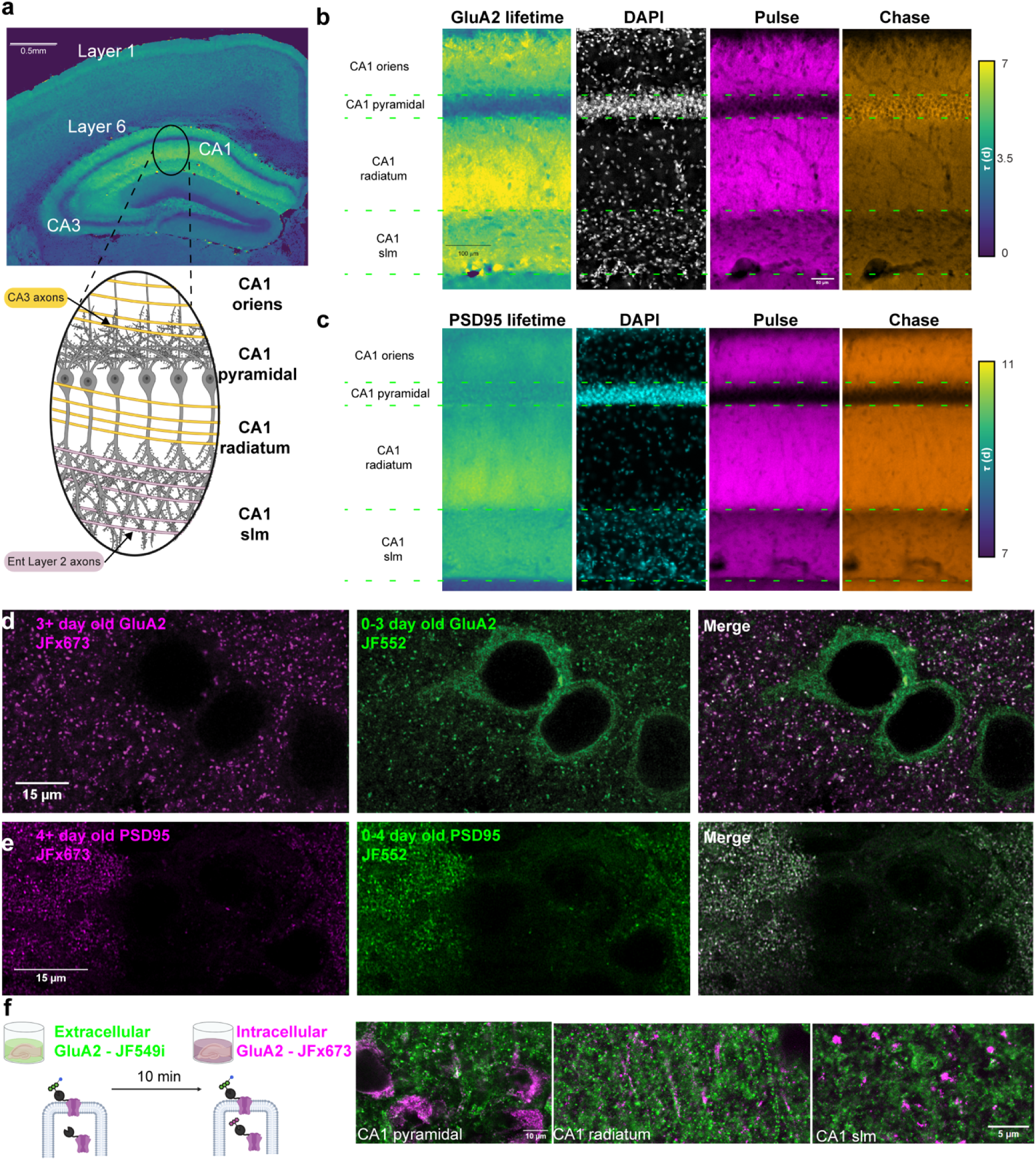
Sub-cellular regulation of synaptic protein turnover. **a, Top**, image of GluA2-HT lifetime in a coronal section around HC. **Bottom**, schematic description of the orientation of the excitatory pyramidal cells in CA1 of HC (gray) with the major axonal input pathways from CA3 (yellow) and from Entorhinal cortex layer 2 cells (pink) **occupying** different layers of the HC. Scale bar is 0.5mm. **b**, Left, GluA2-HT lifetime in days (zoom in; scale bar 0.1mm). 2^nd^ panel, DAPI image showing the transitions of the different HC CA1 lamina. 3^rd^ panel, *Pulse*, 4^th^ panel, *Chase*. We see close to 0 lifetime around the somas, a gradient of lifetimes from the soma into the proximal dendrites in *oriens (so)* and *radiatum (sr)* and a reduction and flat lifetime in CA1 *slm*. The last observation suggests subcellular control of synaptic protein turnover. **c**, same as (b) for PSD95-HT showing very similar turnover trends except for the cell body layer. **d-e**, Differences between GluA2-HT (d) and PSD95-HT (e) in subcellular localization of newly synthesized protein. While 3–4-day old protein (left panels – magenta) is mostly synaptically localized in both GluA2-HT (d) and PSD95-HT (e), newly synthesized GluA2 (green; middle panel in d) is enriched around the soma (in the cytosol while excluded from the nucleus; **Supplementary Movie 2**) while PSD95-HT is not (green; middle panel in e). Merged view in the left panels. **f**, Top: Illustration of an acute live slice experiment that separates two pools of currently expressed GluA2-HT receptors. An extracellular pool is first labeled with a cell-impermeable ligand (JF_549_i-HTL - Green) followed by labeling with a cell permeable one (JF_549_i-HTL - Green). Both pools are seen in somas (2^nd^ panel from top) and both *sr* and *slm* dendrites (3^rd^ and 4^th^ panels).

To explore the possibility of intracellular pools of GluA2, we separated GluA2-HT intracellular and surface pools using a modified pulse-chase paradigm, made possible by the fact that the HT was inserted on the extracellular side of the receptor (**Methods; Fig. 4f**). In freshly cut hippocampal brain slices from GluA2-HT mice, we used a cell-impermeable pulse dye-ligand, JF_549_i-HTL^55–57^, and thus labeled surface receptors only (**JF_549_i; green in Fig. 4f**). After 10 min we used a chase dye that is cell-permeable, and therefore in addition binds intracellular GluA2-HT (**JFX_673_; magenta in Fig. 4f**). CA1 somata were dominated by intracellular GluA2-HT (**Fig. 4f – left panel).** Additionally, both proximal and distal dendrites contained intracellular pools of GluA2-HT (**Fig. 4f – middle and right panels, respectively)**. These results revealed an intracellular (reserve) pool of AMPA receptors^58–62^ throughout CA1 pyramidal neurons (similar reserve pools were observed in the neocortex; **Supplementary Movie 2**).

Together, these results demonstrate the ability of DELTA to identify rich features of subcellular localization and local turnover rates of synaptic proteins.

## Discussion

We developed a robust pulse–chase method, DELTA, to measure turnover for endogenous proteins tagged with HaloTag (HT) at high temporal and spatial precision. We identified the most suitable fluorescent ligands from existing JF dyes and identified JF_552_-HTL, JF_669_-HTL, and its improved variant JFX_673_-HTL^31–33,36^ as the best for this method.

We measured the lifetime of the major excitatory synaptic scaffold protein PSD95 over a range of spatial scales from brain regions to single synapses. PSD95-HT lifetimes ranged from 11–14 days, depending on the brain region (**Fig. 2f, g**), comparable to estimates from metabolic labeling and MS (**Extended Data Fig. 9h,i**), but considerably longer than previous measurements using HT fusions and pulse-only dye-ligand administration^28^. We found that PSD95-HT was destabilized by enriched experience, with middle cortical layers and CA1 as major loci of plasticity.

Using new knock-in mice with HT fused to the extracellular side of GluA2, we found that learning-induced synaptic turnover was most pronounced in the hippocampus, whereas enriched environment caused more widespread changes in turnover. These changes encompassed the entire CA1 hippocampal region, involving many cells. Previous studies have reported small ensembles of memory-related neurons labeled using immediate early gene activation^63,64^. Our results suggest that these small ensembles may represent a subset of highly plastic cells during learning, both locally and brain wide.

We also found evidence for subcellular control of synaptic protein turnover for both GluA2-HT and PSD95-HT, as turnover in the CA1 subregion of the hippocampus was regulated at the level of dendritic compartments (**Fig. 4a-c**). This dendritic specificity was surprisingly similar for GluA2 and PSD95, suggesting coordination of turnover across proteins in a complex^65^. The main difference between GluA2 and PSD95 was seen at the soma, where very little PSD95 protein was observed (**Fig. 4c,e**). These data imply that either PSD95 is rapidly transported into the dendrites after synthesis in the soma and / or is translated in dendrites^66^.

DELTA complements MS-based methods for measuring protein lifetime. MS measures thousands of proteins in parallel but provides only low spatial resolution and sensitivity. For example, EE has been shown to modulate turnover of approximately 30 synaptic proteins^12^ DELTA only measures the lifetime of one HT fusion protein at a time but does so on scales ranging from millimeters to less than 100 nanometers by leveraging modern histology and microscopy methods. Localizing plasticity-related brain regions using DELTA, followed by proteome-wide measurements using MS in these brain regions, could provide comprehensive views of protein turnover in key plasticity loci. DELTA requires genetic manipulation, which itself may influence the target protein’s lifetime, and ligand-dye saturation tests for each new HT fusion. However, DELTA is easily combined with other imaging-based molecular methods, such as fluorescence *in situ* hybridization (FISH)^67^ or immunofluorescence, to provide additional molecular context (**Fig. 1d; Extended Data Fig. 8d-f**). DELTA also has several advantages over pulse-only methods, such as higher SNR, lower variability, and the ability to correct for changes in total protein (**Extended Data Fig. 2f-i, 9e-j; Supplemental Text**).

The ability to perform pulse–chase experiments to study protein turnover *in vivo* will enable experiments that could lead to new insights into the molecular mechanisms of synaptic plasticity and learning. Beyond measuring lifetime for other proteins and in other subcellular compartments, DELTA can be combined with the expression of indicators or actuators of neural activity to examine the relationship between neural activity and protein turnover. This could also be done in both healthy mice and animal models of brain disorders. Furthermore, DELTA is not limited to studies of the brain (**Extended Data Fig. 7**). Overall, DELTA could help identify organ-wide changes involved in adaptive and maladaptive processes in an unbiased, high-throughput, and high-resolution manner.

## Methods

### Materials

JF-HTLs can be requested at HHMI’s Open Chemistry^34^ initiative (dyes.janelia.org) or purchased from Promega. Purified HT protein (#G4491) was from Promega. Pluronic F-127 (#P3000MP), anhydrous dimethylsulfoxide (DMSO; #D12345), *N,N,N*′*,N*′-tetramethylethylenediamine (TEMED; #15524010), ammonium persulfate (APS; #AAJ7632209), and acryloyl-X succinimidyl ester (AcX-SE; #A20770) were from ThermoFisher Scientific. Acrylamide (#1610140) and bis-acrylamide (#1610142) were from Bio-Rad (Hercules, CA). To prepare sodium acrylate for expansion microscopy, acrylic acid (TCI #A0141) was neutralized with sodium hydroxide^68^. 4-Hydroxy-TEMPO (4HT; #176141) was from Sigma (Saint Louis, MO).

### Animals

All procedures were carried out according to protocols approved by the Janelia Institutional Animal Care and Use Committee (IACUC). Wild-type mice (C57Bl/6J RRID:IMSR_JAX:000664; both male and female) were housed in a 12:12 reverse light:dark cycle with *ad libitum* access to food and water until they recovered from a headbar surgery. MeCP2-HT mice^69^, PSD95-HT mice^46^, and GluA2-HT mice (this work) of either sex were used. No comparisons were made between males and females for MeCP2-HT knock-in mice as it is an X-linked gene.

### Dye clearance

Cranial windows were made over the anterior lateral motor cortex or primary visual cortex (centered on - 2.5 mm lateral, + 0.5 mm anterior from lambda) and a headbar was attached posterior to the window^70^. Mice were anesthetized with 1% isoflurane and kept at 37°C on a heating pad. The optical setup^71^ is based on a custom wide-field fluorescence microscope (Thorlabs) and a sCMOS camera (Hamamatsu Orca Flash 4.0 v3). Details of illumination, objectives, and filters are presented in **Supplementary Table 1**. A baseline image was acquired before dye injection. Both the baseline and subsequent timepoints were acquired as movies of 40-50 frames at 5 Hz. This was done to reduce the chances of saturating pixels on the camera at the peak on the injection while still being above electrical noise under baseline conditions. After the estimation of the baseline, the animal was removed from head fixation and injected with dye. After dye injection, it was quickly returned to head fixation; the field of view was compared to a baseline image under room lights and then imaged for up to four hours in intervals of 2 minutes to 20 minutes to cover the fast decay part of the dye clearance. The animal was then recovered and reimaged for up to 24 h at 4-12 h intervals.

The images were analyzed using a custom Matlab script. Briefly, each movie was averaged, and a manual ROI was defined (blue circle in **Extended Data Fig. 1b**). The pixel-averaged mean was fitted to a double exponential: *F* = *a* * *e*^-1/*τ*^_l_ +*b* * *e*^-1/^*^τ^*_2_ + *c*. Population averages were computed by binning all 11 experiments together. In cases where the injection phase was captured, an offset was used to start the fitting from the highest time point.

### Carotid artery dye infusion

To measure the infusion kinetics of JF dyes, we needed a way to control the rate of infusion while imaging dye accumulation in the brain through a cranial window (**Extended Data Fig. 5**). We used two C57Bl/6J mice with carotid artery vascular catheterization (from Charles Rivers surgical service). They were flushed with saline and 50% sucrose lock solution every 7 days. A cranial window was performed 7 days after catheterization as described above. 7 days after the cranial window surgery, a pump with JF dye was connected to the carotid artery line. The initial pump rate was 20 µL/min followed by 40 µL/min and 80 µl/min. The imaging conditions were the same as for the dye clearance experiments (0.2 s exposure) but imaged continuously for each injection speed. Data were time averaged at one second intervals (5 timepoints) and a manual ROI was drawn. The time-averaged fluorescence values were trimmed to the times the pump was running and fit with a linear fit (including intercept). The values reported are the slopes of the fit.

### Modeling dye clearance effects on saturation

Given our results on dye infusion (**Supplementary Text; Extended Data Fig. 5e**), we tested the hypothesis that given a fixed amount of dye to inject, faster injections would lead to more dye captured in the brain. We used a model where a single position in space (x position 1) is a blood vessel from which dye can enter the brain and diffuse in 1D simulated using a time march algorithm (Finite difference method). To compare the effect of injection speed alone, in each simulation, we used the same total amount of dye injected, and the same amount of dye was cleared from the vessel (1AU/dt). Simulations had different injection rates (2-10 AU/dt) and the width of the dye distribution was measured as the width at 90% of the known saturation value.

### Modeling protein turnover measurements with DELTA

Pulse-chase experiments were modeled using the SimBiology package in Matlab. Lifetimes of the target HT-protein synthesis were simulated. Each new protein could either degrade (at the same rate of synthesis modeled) or attach to a HTL dye molecule (pulse or chase, dye collectively). The dye binding kinetics were set to be instantaneous, as they are several orders of magnitude higher than those for synthesis/degradation. Binding kinetics were identical for both the pulse and chase. The degradation rates of the HT protein with dye complexes were the same as for the protein alone.

The dye injection kinetics were neglected (*i.e.*, assume instantaneous injection). Dye clearance kinetics were modeled with two compartments (cytosol and lipids) with clearance from the cytosol compartment. Multiple compartments were required to account for the multiexponential decay measured *in vivo*. The cytosol compartment was set to have a volume of 1 mL; the equilibrium constant between the compartments, the volume of the lipids compartment and the degradation time constant were fitted using the JF-dyes clearance data (**Extended Data Fig. 1,2**). This fitting was done in the absence of HT-protein.

For evaluating the effects of changing total protein, we assumed ideal measurement conditions (single exponential decay, no dye clearance kinetics or noise). We calculated the difference between the estimated lifetime of a protein (τ = 3 days) under same total protein levels vs. changing the total amount of protein at the end of the 3 days (**Extended Data Fig. 2i**).

### Dye solubility

JF dye ligands were prepared in the following manner. (i) Captisol formulation was made with 300 mg Captisol® (β-Cyclodextrin Sulfobutyl Ethers, sodium salts; NC-04A-180186, Cydex Pharmaceuticals) dissolved in 1 mL of water (Molecular Biology Grade Water, Corning) to make a 30% solution. 100 µl of this solution was added to 100 nmol of dry JF dye. (ii) A Captisol + Pluronic formulation was made by mixing a 30% Captisol solution with Pluronic F-127 (P3000MP, Invitrogen) in an 80:20 ratio. 100 µl of the prepared solution was added to 100 nmol of dry JF dye. (iii) The DMSO + Pluronic formulation was made with DMSO (D2650, Sigma Aldrich), Pluronic F-127, and saline (0.9 % sodium chloride) mixed in a 20:20:60 ratio. 100 µL from the prepared solution was added to 100 nmol of dry JF dye. (iv) The DMSO formulation was made by adding 100 µl of DMSO to 100 nmol of dry JF dye to bring it to a final concentration of 1 mM.

The prepared JF dye formulations were briefly vortexed followed by bath sonication (Branson 1200 model) for 5 minutes. The dye solutions were placed on the agitator for 72 h to ensure solubilization. Absorbance measurements of dye solutions were performed using a spectrometer (Cary 100 UV-Vis, Agilent Technologies), and the final concentration was determined from known extinction coefficients of JF dyes (**Extended Data Fig. 7e-h**)^31–33,36^.

### Evaluation of JF-HaloTag ligands *in vivo*

To evaluate the ability of different JF HTL dyes to saturate HaloTag proteins in the brain we expressed HT-EGFP (**Extended Data Figure 3,4**). Virus delivery for sparse expression and dye delivery were achieved using retro-orbital (RO) injections^72^. Dye preparation^73^, dye injections^74^, and histological preparations^75^ have been described. A virus to express HT-EGFP was prepared using a PHP.eB AAV capsid^76^ with a synapsin promoter. 100 μL of virus (titer of 4e11 GC/mL) was injected RO. An mKate-HT was used to avoid cross-talk between the GFP and the JF_525_-HTL dye. RO dye injections (pulse) were performed 3-5 weeks after the viral injection. Dyes were prepared by dissolving 100 nmol of dye in 20 µl DMSO followed by 20 µL of 20% Pluronic F-127 in DMSO and 60 µl of PBS, except when otherwise stated (Supplemental Table 2). Twenty-four hours after dye injection, animals were perfused with 50 mL of 4% PFA in 0.1 M sodium phosphate, pH 7.4 (PB) and 50 nmol of orthogonal dye. The brains were post-fixed in 4% PFA in 0.1 M PB overnight at 4°C and washed three times in PBS for 15 minutes. 100 µm coronal sections were cut and floated in 24-well plates followed by 4 h of DAPI staining (0.6 µM in PBS) and washed 3 times for 15 minutes in PBS. For animals with visible fluorescence from *in vivo* injection, every fourth slice was mounted for imaging; for animals where no fluorescence was observed, every 24^th^ slice was mounted and imaged. See Supplementary Table 2 for details of the animals in the section.

### Immunofluorescence

For MeCP2-HT mice we performed Immunofluorescence (IF) staining to distinguish cell types in histological sections (**Fig. 1d; Extended Data Fig. 8d-f**). Brains were sectioned coronally, and sections were blocked in PBS with 2% BSA and 0.1% Triton X-100 for 1h at room temperature. Primary antibodies (Rabbit Anti-NeuN RRID: AB_10807945 Millipore Cat# ABN78; Rabbit Anti-Iba1 RRID: AB_2636859 Abcam Cat# ab178846; Rabbit Anti-SOX10 RRID: AB_2721184 Abcam Cat# ab180862) were applied at a concentration of 1:250 n in the same buffer overnight at 4°C. The sections were washed three times for 15 minutes. The secondary antibody was a Goat anti-Rabbit AF488 (RRID: AB_143165) used at 1:500 overnight at 4°C. After three more 15-minute washes, the slices were stained with DAPI as described^75^ and mounted. For PSD95 immunofluorescence (**Extended Data Fig. 9c**), we used a validated anti-PSD95 antibody (NeuroMAB, RRID: AB_10807979) with a Goat anti-mouse CF633 secondary (RRID: AB_10854245). For validation of GluA2-HT (**Extended Data Fig. 10c**), we used another monoclonal antibody from NeuroMAB (RRID: AB_2232661) with the same secondary.

### Expansion Microscopy

Coronal sections (**Fig. 2k**; **Fig. 4d,e**; **Extended Data Fig. 9c; Extended Data Fig. 10c**) were anchored with AcX (0.033 mg/mL; 1:300 from 10 mg/mL stock in DMSO) in PBS for one hour. Sections were then transferred to a gelation solution (2.7 M acrylamide, 0.2 M sodium acrylate, 200 µg/mL bis, PBS (1×), 2 mg/mL APS, 2 mg/mL TEMED, and 20 µg/mL 4HT) and incubated on ice, shaking, for 30 minutes before being mounted in gelation chambers and moved to 37 °C for one hour. Excess gel was removed, and the tissue was recovered in pure water. With three one-hour washes in pure water, the slices expanded approximately 2-fold without disruption or cracks. These sections were moved to a 6-well glass bottom plate for microscopy (Cellvis #P06-1.5H-N, Mountain View, CA). To flatten and immobilize the sections, 4 dabs of silicon grease were applied around each section and a 22x22 mm square #2 coverslip (Corning #2855-22, Corning, NY) was pressed from above. If needed, poly-L-lysine 0.2 mg/mL (Sigma #P1524-25MG) with Photo-Flo 200 (1:500 from stock; #74257 Electron Microscopy Sciences, Hatfield, PA) were applied to the bottom of the well before the sections are placed to better immobilize the gels.

### Brain-wide imaging

Imaging of entire sections at high resolution was performed on a confocal slide scanner consisting of a TissueFAXS 200 slide feeder (Tissuegnostics, Germany) and a SpectraX light engine (Lumencor). Illumination light (wavelength, power, (excitation filters center / width): (1) 395 nm, 400 mW (395 nm / 25 nm); (2) 475nm, 480mW (475 nm / 34 nm); (3) 400mM (585 nm / 35 nm); (4) 619 nm, 629 mW (635 nm / 22 nm) was delivered by a lightguide to a Crest X-Light V2 confocal spinning disc microscope (Crestoptics; 60 μm pinhole spinning disk) with the following dichroics (Chroma Technology, T425lpxr, T495lpxt, T600lpxr, T660lpxr and emission filters (ET460 nm / 50 nm, ET525 nm / 50 nm, ET625 nm / 30 nm, ET700 nm / 75 nm). The emission light was collected with Zeiss objectives: EC Plan-Neofluar 10x/0.3 M27 for MeCP2 and PSD95 animals (**Fig. 2c-k; Extended Data Fig. 6,7**) and a Plan-Apochromat 20x/0.8 M27 for virally injected animals (**Extended Data Fig. 3,4**). Detection was with a Zyla 5.5 sCMOS camera (Andor). Image acquisition involved semi-automated tissue detection using multiple autofocusing points per section (5x5 and 3x3 grids for objective acquisitions of 20x and 10x accordingly). For virally transfected animals, three z-planes were imaged with a 7 µm spacing and projected in z. For MeCP2 and PSD95 animals, a single plane was imaged.

### High- and Super-resolution imaging

To segment individual nuclei of MeCP2-HT-expressing cells (**Fig. 1d-f**; **Extended Data Fig. 8**) or single synapses expressing PSD95-HT (**Fig. 2i**), higher resolution imaging was needed. For MeCP2 imaging we used a Zeiss LSM 880 (Blackwood, NJ) with an EC Plan-Neoflour 40x/1.3NA oil objective and a voxel size of 0.25x0.25x0.5 μm. We acquired 21 z planes in 3 tracks (a track can have more than one detection channel for multiplexing) and 5 channels (Detection [excitation] wavelengths were as follows (all in nm): Track 1 - DAPI: 410-489 [405]- DAPI, JF_612_: 624-668 [594]; Track 2 IF: 493-551 [488], JF_669_: 680-758 [633]; Track 3 JF_552_: 553-588 [561]. For PSD95 imaging we used a Zeiss LSM 980 with Airyscan and a C-Apochromat 40x/1.2 water dipping objective. The voxel size was 57x57x230 nm. We acquired 23 z sections with two channels, 561 nm illumination for JF_552_ and 633 nm illumination for JFX_673_. A full z-stack was acquired for the far-red channel followed by the red channel.

### Image analysis

#### Janelia Flour dye screening

We used *Fraction Pulse* = *Pulse*/(*Pulse+Chase*) a measure of saturation, where (*Pulse+Chase*) is a measure of total protein. *Fraction Pulse* was calculated after conversion to µM dye, where adding the pulse and the chase are a valid operation (same units). *Fraction Pulse* values closer to 1 indicate saturation. Doing a pixel-wise analysis would not be sufficient, as there are background signals that would change depending on the imaged channel and brain region. Here, unlike knock-in animals (GFP-HT expression with the PHP.eb virus), there is variability in the expression pattern across animals that could also affect the background (neuropil) in which the cells reside (**Extended Data Fig. 3**).

We performed analysis on detected cells with local background subtraction. To detect cells we imported downsampled (2x) images with three channels (GFP/*Pulse*/*Chase*) into Ilastik’s pixel classification workflow^77^. The workflow was manually trained to segment cell bodies, dendrites, neuropil signal, and various imaging and tissue artifacts. The pixel probability maps for cells, combined with the raw data, were then imported to a second object-classification workflow in Ilastik. This workflow was used to classify each mask as a GFP-HT-expressing cell or not. A Matlab (Mathworks) script was used to calculate the value of each mask in the three channels and to calculate a local background. The local background estimation excluded pixels that belonged to other non-neuropil trained categories (other cells, dendrites, etc.) from the first Ilastik workflow (**Extended Data Fig. 3c**). For each mask, the local background was subtracted, and the fluorescence value was converted to µM dye concentration using an independent calibration obtained under the same imaging conditions (**Extended Data Fig. 4a**).

#### MeCP2-HaloTag knock-in mice

Nuclei were segmented and categorized into cell types using IF (**Fig. 1c-e; Extended Data Fig. 8d-f**). First, the five imaged channels (DAPI, *Pulse*, *Chase1*, *Chase2*, IF) were each normalized (0 to 1) using the top and bottom 0.3% of the pixel intensity histogram. Second, the mean of all three imaged JF dyes was used to make a 3-channel image (DAPI, mean of all JF dyes, IF). This 3-channel image was used in a pixel classification workflow using Ilastik (ilastik.org). We used training data from all three types of IF (NeuN, Iba1, and SOX10). The resulting nuclei probability maps, together with the 3-channel images, were used by three independent object classification workflows in Ilastik, one for each IF type. The output was a set of masks for IF-positive nuclei for each cell type. We calculated three *Fraction Pulses* from these dual Chase experiments for each segmented nuclei: (1) *Pulse* / *Pulse* + *Chase 1*; (2) *Pulse + Chase 1* / *Pulse* + *Chase 1* + *Chase 2*; (3) *Chase 1* / *Chase 1* + *Chase 2*. These were done after conversion to μM dye units using a calibration curve for each dye making the addition of different dyes a valid operation. We then pooled all 15 measurement (5 animals x 3 *Fraction Pulses*) and used bootstrapping to fit a lifetime with the assumption of a single exponential decay.

#### PSD95-HaloTag mice

In contrast to MeCP2, the diffuse PSD95 expression did not permit segmentation of individual neurons or synapses with standard fluorescence microscopy. We instead performed a pixel-based analysis after 2x down sampling (resulting pixel size, 2.7 × 2.7 µm) (**Fig. 2b-k**). As with MeCP2, we converted our images to 3-channel images (DAPI/*Pulse*/*Chase*). A pixel-classification workflow (Ilastik) was trained to exclude ventricles and artifacts. Each pixel was converted to a lifetime estimate with the assumption of a single exponential decay after conversion to μM dye concentration using calibration curves imaged under the same conditions as was explained above.

#### GluA2-HaloTag mice

GluA2 analysis was similar to PSD95, with slight modifications. A different workflow was trained (Ilastik) to segment regions with signal. Additionally, four animals without HT (negative controls) were imaged, registered to the Allen Brain Reference Atlas^49^ (CCFv3) and their pulse and chase signals were quantified. This negative control was used to define a brain-region specific threshold as 2x the standard deviation above the mean of the negative control average. The threshold was used to exclude brain regions where GluA2-HT signal was not significantly higher than autofluorescence. This was done for both the pulse and the chase channels, causing rejection of 140/1327 possible regions (CCF v3). As CA1 lamina (**Extended Data Fig. 10k**; *so*, *sr*, *slm* and the cell body layer) were not present in our reference atlas (CCFv3) we manually segmented them using the Ground Truth Labeler GUI in Matlab.

#### Single synapse analysis after Expansion Microscopy

After acquisition using the Airyscan detector array, we used Zen Blue software (Zeiss) to process the images (Airyscan processing function in 3D with automated deconvolution strength). The Airyscan-processed images were registered across channels as they were acquired sequentially. The resulting 2-channel image was normalized and used as input to an Ilastik pixel classification pipeline trained to separate synapses from the background. The resulting probability images with the normalized data were used in a second Ilastik object segmentation pipeline (**Fig. 2l**). A local background was subtracted, fluorescence values were converted to μM dye, and lifetimes were estimated.

### Alignment to the Allen Brain Reference Atlas CCFv3

MeCP2, PSD95 and GluA2 lifetime measurements were registered to the Allen Brain Reference Atlas^49^ (CCFv3) using a two-step procedure. First, a downsampled 24-bit RGB image of each section was loaded with QuickNII^78^ and aligned with the 25 µm voxel resolution version. This accounted for the cutting angle, a global scaling factor in the dorsoventral and mediolateral axes, and the anteroposterior location of each section. QuickNII output was used for a manual non-rigid alignment with VisuAlign^79^. The main markers for registration were the edges of the section, the ventricles, and the fiber tracks. The VisuAlign output was another RGB image in which each color was assigned to an Allen CCFv3 id. These images were loaded and interpolated to the original size of each section, allowing the assignment of each MeCP2 nucleus and PSD95/GluA2 pixel to a CCF id. As the PSD95 and GluA2 analysis was pixel based, we excluded pixels belonging to the root or any part of the CCFv3 tree under fiber tracks or ventricular system.

### Statistical analysis

To compare brain regions across animals assigned to different experimental conditions, we employed a Linear Mixed Effects (LME) model. Given that data was acquired from coronal slices, where the same brain regions may appear in varying numbers of slices across different animals, we used pixels as our base unit of measurement. Each pixel was represented as a row in a table with columns corresponding to the animal ID, brain region, and mean lifetime.

We fitted an LME model using this table, applying median filtering to cap the maximum number of repeated measurements at 1,000 per condition (where a condition is defined as the same brain region within the same animal). The LME model included one fixed effect (group assignment: Control, EE, Baseline, New Rule), one random effect (animal ID), and an estimate of residual error.

This approach allowed us to compute the mean and standard error of the mean for each condition in the fixed effects (e.g., Layer 1, Layer 2/3), as well as the mean and standard deviation for the residual error and across conditions in the random effects (e.g., individual animals).

### Mass spectroscopy (MS)-based measurement of turnover

#### C13 lysine heavy isotope feed labeling in mice and MS sample preparation

We followed a previously published protocol for isotope labeling for MS based turnover measurements^80^. Either wild-type or homozygote PSD95-HT mice cortices were rinsed and dissected in solution A (5 mM HEPES pH 7.4, 1 mM MgCl2, 0.5 mM CaCl2, 1 mM DTT, 0.32 M sucrose and protease and phosphatase inhibitor set (Thermo Fisher Scientific, Cat#: 78443_3670527377) on ice. Then the tissues were homogenized with an electronic homogenizer (Glas-Col, Cat# 099C-K54). Homogenates were spun down at 1,400 g for 10 mins (4°C). Set aside the supernatant. We then resuspended the pellets in 20 mL solution A. The diluted homogenates were centrifuged at 710 g for 10 mins (4°C). The pellet is P1. We combined and mixed the supernatant and the saved supernatant as S1. S1 was centrifuged at 13,800 g for 10 mins (4°C). The supernatant is S2. Then we resuspended the pellets (P2) in solution B (6 mM Tris pH 8.1, 0.32 M sucrose, 1 mM EDTA, 1 mM EGTA, 1 mM DTT with protease and phosphatase inhibitors).

Proteins were extracted from P2 using a chloroform-methanol precipitation method. Protein pellets were resuspended in 8 M urea (Thermo Fisher Scientific, Cat # 29700) prepared in 100 mM ammonium bicarbonate solution (Fluka, Cat # 09830). The samples were reduced with 5 mM Tris(2-carboxyethyl)phosphine (TCEP, Sigma-Aldrich, Cat # C4706; vortexed for 1 hour at RT), alkylated in the dark with 10 mM iodoacetamide (IAA, Sigma-Aldrich, Cat # I1149; 20 min at RT), diluted with 100 mM ABC, and quenched with 25 mM TCEP. Samples were diluted with 100 mM ammonium bicarbonate solution, and digested with Trypsin (1:50, Promega, Cat # V5280) for overnight incubation at 37°C with intensive agitation. The next day, the reaction was quenched by adding 1% trifluoroacetic acid (TFA, Fisher Scientific, O4902-100). Samples were desalted using Peptide Desalting Spin Columns (Thermo Fisher Scientific, Cat # 89882). All samples were vacuum centrifuged to dry. We further used a high pH reverse-phase peptide fractionation kit (Thermo Fisher Scientific, Cat# 84868) to get eight fractions (5.0%, 7.5%, 10.0%, 12.5%, 15.0%, 17.5%, 20.0% and 50.0% of ACN in 0.1% triethylamine solution). All fractions were vacuum centrifuged to dry. The high pH peptide fractions were directly loaded into the autosampler for MS analysis without further desalting.

#### Tandem mass spectrometry

Three micrograms of each sample were auto-sampler loaded with a Thermo Vanquish Neo UHPLC system onto a PepMap™ Neo Trap Cartridge (Thermo Fisher Scientific, Cat#: 174500, diameter, 300 µm, length, 5 mm, particle size, 5 µm, pore size, 100 Å, stationary phase, C18) coupled to a nanoViper analytical column (Thermo Fisher Scientific, Cat#: 164570, diameter, 0.075 mm, length, 500 mm, particle size, 3 µm, pore size, 100 Å, stationary phase, C18) with stainless steel emitter tip assembled on the Nanospray Flex Ion Source with a spray voltage of 2000 V. An Orbitrap Ascend (Thermo Fisher Scientific) was used to acquire all the MS spectral data. Buffer A contained 99.9% H_2_O and 0.1% FA, and buffer B contained 80.0% ACN, 19.9% H_2_O with 0.1% FA. For each fraction, the chromatographic run was for 2 hours in total with the following profile: 0-8% for 6, 8% for 64, 24% for 20, 36% for 10, 55% for 10, 95% for 10 and again 95% for 6 mins receptively.

We used Orbitrap HCD-MS2 method for these experiments. Briefly, ion transfer tube temp = 275°C, Easy-IC internal mass calibration, default charge state = 2 and cycle time = 3 s. Detector type set to Orbitrap, with 60K resolution, with wide quad isolation, mass range = normal, scan range = 375-1500 m/z, max injection time mode = Auto, AGC target = Standard, microscans = 1, S-lens RF level = 60, without source fragmentation, and datatype = Profile. MIPS was set as on, included charge states = 2-7 (reject unassigned). Dynamic exclusion enabled with n = 1 for 60s exclusion duration at 10 ppm for high and low with Exclude Isotopes. Isolation Mode = Quadrupole, isolation window = 1.6, isolation Offset = Off, active type = HCD, collision energy mode = Fixed, HCD collision energy type = Normalized, HCD collision energy = 25%, detector type = Orbitrap, orbitrap resolution = 15K, mass range = Normal, scan range mode = Auto, max injection time mode = Auto, AGC target = Standard, Microscans = 1, data type = Centroid.

#### MS data analysis and quantification

Protein identification/quantification and analysis were performed with Integrated Proteomics Pipeline - IP2 (Bruker, Madison, WI. http://www.integratedproteomics.com/) using ProLuCID^81,82^, DTASelect2^83,84^, Census and Quantitative Analysis. Spectrum raw files were extracted into MS1, MS2 files using RawConverter (http://fields.scripps.edu/downloads.php). The tandem mass spectra (raw files from the same sample were searched together) were searched against UniProt mouse (downloaded on 07-29-2023) protein databases^85^ and matched to sequences using the ProLuCID/SEQUEST algorithm (ProLuCID version 3.1) with 50 ppm peptide mass tolerance for precursor ions and 600 ppm for fragment ions. The search space included all fully and half-tryptic peptide candidates within the mass tolerance window with no-miscleavage constraint, assembled, and filtered with DTASelect2 through IP2. To estimate protein probabilities and false-discovery rates (FDR) accurately, we used a target/decoy database containing the reversed sequences of all the proteins appended to the target database^85^. Each protein identified was required to have a minimum of one peptide of minimal length of six amino acid residues. After the peptide/spectrum matches were filtered, we estimated that the protein FDRs were ≤ 1% for each sample analysis. Resulting protein lists include subset proteins to allow for consideration of all possible protein isoforms implicated by at least three given peptides identified from the complex protein mixtures. Then, we used Census and Quantitative Analysis in IP2 for protein quantification. Static modification: 57.02146 C for carbamidomethylation. Differential modification: 42.0106 for acetylation at N-terminals. Mass shift for heavy lysine, 6.0201. Quantification was performed by the built-in module in IP2. For quantification, only peptides containing the amino acid lysine (K) were selected. For the estimation of protein half-life, H/L values were analyzed as previously described^12,80^. Half-life values were divided by ln(2) to calculate mean lifetime (τ).

### GluA2-HaloTag knock-in mouse generation and validation

The GluA2 Halo knock-in line was generated by Crispr/cas9 pronuclear microinjection. The Halo gene was inserted after the signal peptide of the GluA2 gene and flanked by two linkers, GGGGSGGGS at the 5’ end and GGGGSGGGSGGGGSGGGS at the 3’end. The homologous arms are 332 bp and 655 bp respectively. The knock-in DNA was co-injected with a gRNA (5’- TCTTCTAACAGCATACAGATAGG-3’) and Cas9 protein (Fisher Sci. Cat A36498) into B6D2F1/J mouse one cell embryos. F2’s were used in this study from both sexes. A founder with germline transmission is being backcrossed to a C57BL/6J background and will be donated to Jackson Laboratories for dissemination.

For brain protein level measurements (**Extended Data Fig. 10b),** hemibrains were freshly dissected and homogenized in 1 mL of ice-cold homogenization solution (320mM sucrose, 10 mM HEPES pH 7.4, 1 mM EDTA, 5 mM Na pyrophosphate, 1 mM Na_3_VO_4_, 200nM okadaic acid, protease inhibitor cocktail (PIC, Roche). The resulting homogenate was centrifuged at 800 x g for ten minutes at 4°C to obtain the P1 (nuclear, pellet) and S1 (supernatant) fractions. We then centrifuged the S1 fraction at 17,000 x g for 20 minutes at 4°C to obtain the P2 (membrane, pellet) and S2 (cytosol, supernatant) fractions. Mouse brain P2 protein fractions were separated by SDS-PAGE using 6% gels and proteins transferred onto nitrocellulose membranes for western blot analysis. Western blots were imaged using the Licor Odyssey M imager. The following antibodies were used for detection: JH6773 (homemade) anti-GluA2 rabbit polyclonal antibody, anti-HaloTag monoclonal antibody (Promega #G921A), and anti-PSD95 monoclonal antibody K28/74R (Addgene #184184 see^86^). JH6773 is a GluA2 N-terminal antibody generated using CDYDDSLVSKFIERWSTLE peptide (amino acids 264-281). For protein quantification, three wild-type and three homozygous GluA2-HT mouse brain samples were run in duplicate, quantified using Licor Image Studio software, and the averages for each duplicate were taken to measure expression levels using PSD95 levels as a normalization signal.

### Brain slice preparation and electrophysiology

For the validation and investigation of GluA2-HT mice (**Fig. 4f, Extended Data Fig. 10e-h**), transverse hippocampal slices (300 µm) were prepared from 3-6 months old wild type or GluA2-HT knock-in mice. The brain was removed rapidly and mounted in a near-horizontal plane for slice preparation (vibrating tissue slicer Leica VT 1200S, Leica Microsystems, Wetzlar, Germany). Slices were prepared in ice-cold sucrose-based solution containing (in mM): 75 Sucrose, 75 NaCl, 2.5 KCl, 25 NaHCO_3_, 1.25 NaH_2_PO_4_, 7 MgCl_2_, 0.5 CaCl_2_, 25 Dextrose. Slices were then transferred to an incubation chamber filled with oxygenated ACSF containing (in mM): 125 NaCl, 2.5 KCl, 25 NaHCO_3_, 1.25 NaH_2_PO_4_, 1 MgCl_2_, 2 CaCl_2_, 25 Dextrose. After half an hour of recovery at 35°C, the slice chamber was maintained at room temperature. All experiments were performed in the presence of GABA_A_ and GABA_B_ receptor antagonists SR95531 (4 μM) and CGP52432 (1 μM).

Whole-cell current-clamp recordings were obtained with a Multiclamp 700B amplifier (Molecular Device, San Jose, CA), using bridge balance and electrode-capacitance compensation. Patch-clamp electrodes were pulled from borosilicate glass (1.5 mm outer diameter) and filled with intracellular solution containing (mM): 120 K-gluconate, 20 KCl, 10 Na_2_phosphocreatine, 10 HEPES, 2 MgATP, 0.3 Na2GTP, 0.1% Biocytin. Electrode resistance in the bath was 3-5 MΩ and series resistance during the recordings ranged was 15-30 MΩ. Electrophysiological traces were low-pass-filtered with a cut-off frequency of 5 kHz and digitized at 20 kHz via a USB-6343 board (National Instruments) under the control of WaveSurfer software (https://wavesurfer.janelia.org/; Janelia Scientific Computing).

EPSP amplitude was monitored while stimulating extracellularly at 0.05 Hz with concentric bipolar electrodes (FHC, Bowdoin, ME) with A365 stimulus isolator (World Precision Instruments, Sarasota, FL). Stimulating electrodes were positioned in the distal *stratum radiatum*, closer to the *stratum lacunosum-moleculare* than the *stratum pyramidale*, at least 200 μm away from the recorded cell body. LTP was induced using a theta-burst protocol in which a single burst of EPSPs (five stimuli, 100 Hz) was delivered five times at 5 Hz. The theta stimulus was repeated 3 times, separated by 30 s. Membrane potential s.d. was estimated by using 2-5 min recordings at -65 mV without synaptic stimulation. Analysis of electrophysiology data was performed using Clampfit 11.3 (Molecular Device, San Jose, CA) and statistical tests were performed using Prism 10. (GraphPad Software, Boston, MA).

Separate brain slices were used to localize the intracellular and extracellular pools of GluA2-HT receptors (**Fig. 4f**). A cell-impermeable version of JF_549_ (JF_549i_-HTL)^32^ was first dissolved in DMSO to a concentration of 1 mM. A working solution concentration of 1 μM was applied for 10 min in ACSF. After 10 min a cell-permeable dye was added (1 μM JFx_673_-from 1 mM stock dye in DMSO). Following another 10 min, 3 × 5min washes were done in ACSF before the slices were fixed for 30 min at room temperature with 4% PFA in 0.1 M sodium phosphate, pH 7.4. Slices were washed 3×15 min in 1× PBS and expanded 2-fold as described below. These short incubation periods meant the dyes did not penetrate the whole 300 μm of the slice. Images were taken from depths where saturation of the cell-impermeable dye was observed (5-50 μm deep) as seen by the separation of extracellular staining and intracellular staining whereas deeper in the tissue the cell-permeable dye was seen in both compartments.

### Behavior protocols for GluA2-HaloTag

#### Headbar implant for behavior in virtual reality

We followed a previous protocol^87^ with slight modifications. Mice were anesthetized with 1.5–2% isoflurane during the time of the procedure. We exposed the skull, using the following coordinates (in mm): 1.8 posterior relative to bregma and 2.0 lateral relative to the midline. Once we localized the A/P and M/L coordinates we drew a dot on the skull. We centered the headbar around this point and we glued the titanium headbar (custom design) with dental cement in the skull. After surgery, we allowed animals to recover for a minimum of 3 days before their water restriction protocol.

#### Water restriction and handling

After surgery mice were single housed and underwent water restriction (typically 1–1.5 ml/day depending on their weight) for 3–5 days before initiating training. During water restriction mice were gently handled and daily water was delivered with a syringe directly to the mouse’s mouth. In addition, we provided the mice with snacks; sunflower seeds, cocoa krispies, or fruit loops. At the end of the session, we provided the animals with a yogurt drop. These handling sessions lasted at least 10 min.

#### Virtual reality arena

Head-fixed mice ran atop a foam ball^88^ in a VR environment with screens on each side and in front of their heads (**Fig. 3a**). Mice were first acclimatized to the ball without visual stimuli (two daily sessions, 45 minutes), while water was delivered at random intervals. Mice were then trained to lick when one of two stimuli appeared in random order in the VR corridor every 30-70 cm of running selected from a uniform distribution. Reward probability was 50%. During the first 2-3 sessions rewards were delivered automatically when the mouse ran through both cues. In later sessions water rewards were delivered only in response to answer licks at the cues. After each behavioral session water was supplemented to a total of 0.5-1 ml.

### Data and Code availability

Both metadata and raw data are available through the Open Science Foundation <u>project</u> associated with this paper. The protocols are available as a collection on <u>protocols.io</u>. The code used for the analysis is available on <u>GitHub</u>. We will deposit the raw MS data into public databases upon acceptance.

## Acknowledgments

We thank Caiying Guo for the generation of the GluA2-HT lines; Benjamin Foster, Monique Copeland, Amy Hu, Susan Michael for histology; Michael DeSantis and Damien Alcor for help with imaging; Gillian Harris, Mariam Rose, Sarah Lindo for help with animal husbandry and surgery; Alina Gutu, Kim Ritola, Hyuh Ah Yi for viral reagents; Ariana Tkachuk, Anastasia Osowski, Katie Holland for help with JF HTL dyes; Eric Schreiter for the GFP-HT virus; James Liu for the MeCP2-HT mouse, and Seth Grant for the PSD95-HT mouse. We thank Amrita Singh for assistance with dye clearance imaging. We thank the Proteomics Core at Northwestern University for their help with the MS measurements. We thank HHMI for funding the research. EFF is supported by a CZI Collaborative Pairs Pilot Project Awards (Cycle 2; Phase 1). EFF also acknowledges the support of the SFB1286, Göttingen, Germany. JNS acknowledges support from NIH grant S10 OD032464. TT received funding from the European Research Council (ERC) under the European Union’s Horizon 2020 research and innovation program (‘MolDynForSyn,’ grant agreement No. 945700).

## Extended Data Figures

**Extended Data Figure 1.**
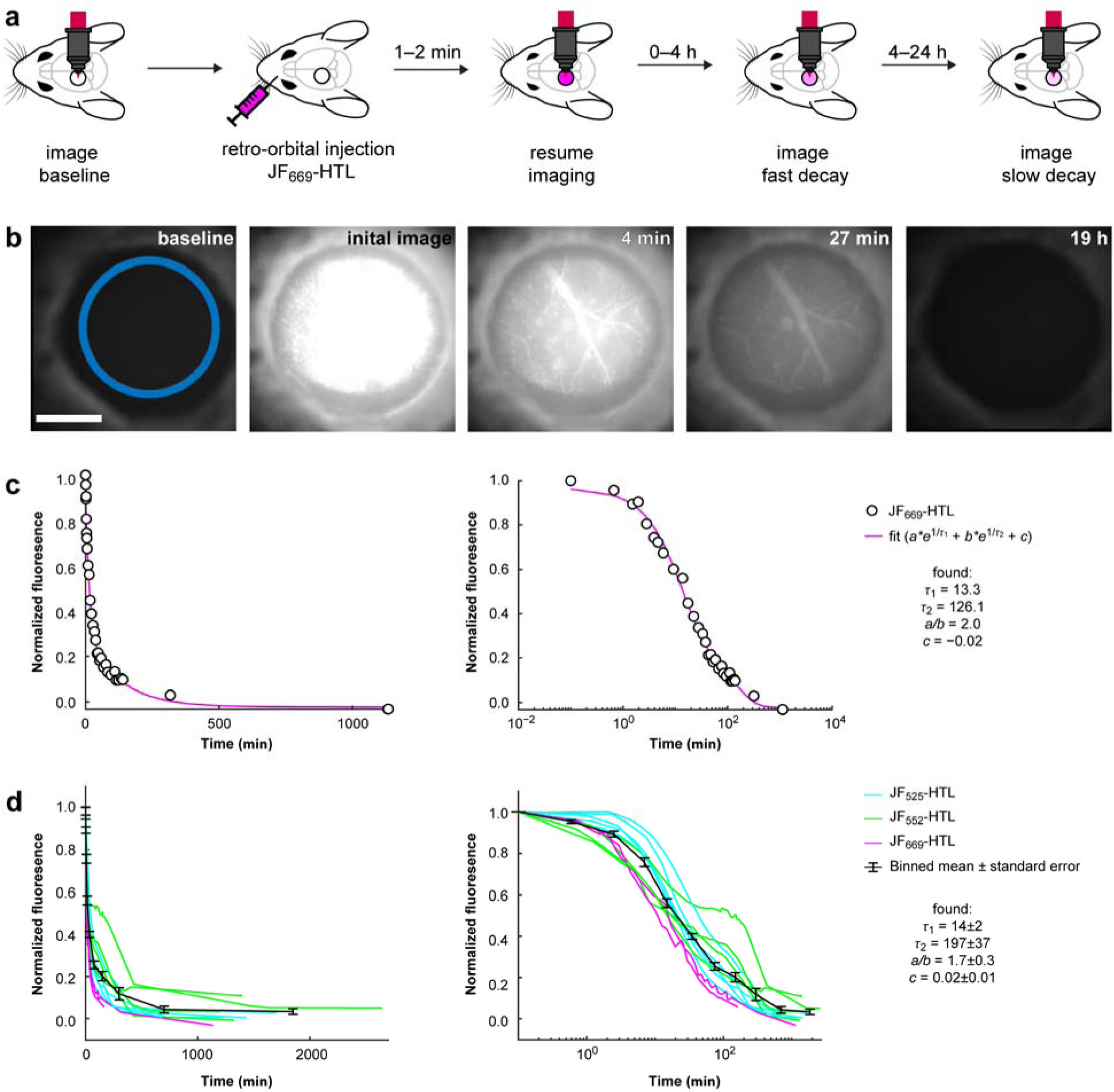
HaloTag ligand dyes are cleared from the brain within hours. **a,** Experimental procedure to measure dye clearance. The rise in fluorescence is not captured, as the animal is not imaged during the injection. **b,** Example images for a JF_669_-HTL injection. The blue circle represents the region for which fluorescence was quantified in (c). Scale, 2 mm. **c,** Linear (left) and log (right) fluorescence (peak normalized and baseline subtracted) as JF_669_-HTL is cleared from the brain. Double exponential fit in red; r_1_ and r_2_ are in minutes. **d,** Similar data for 12 experiments with **3** dyes (JF_525_-HTL in green, JF_552_-HTL in red, and JF_669_-HTL in magenta). The black trace is a binned average, with standard error for all **12** experiments. The numbers in (d) represent the mean ± standard error of double exponential fits for the 12 experiments.

**Extended Data Figure 2.**
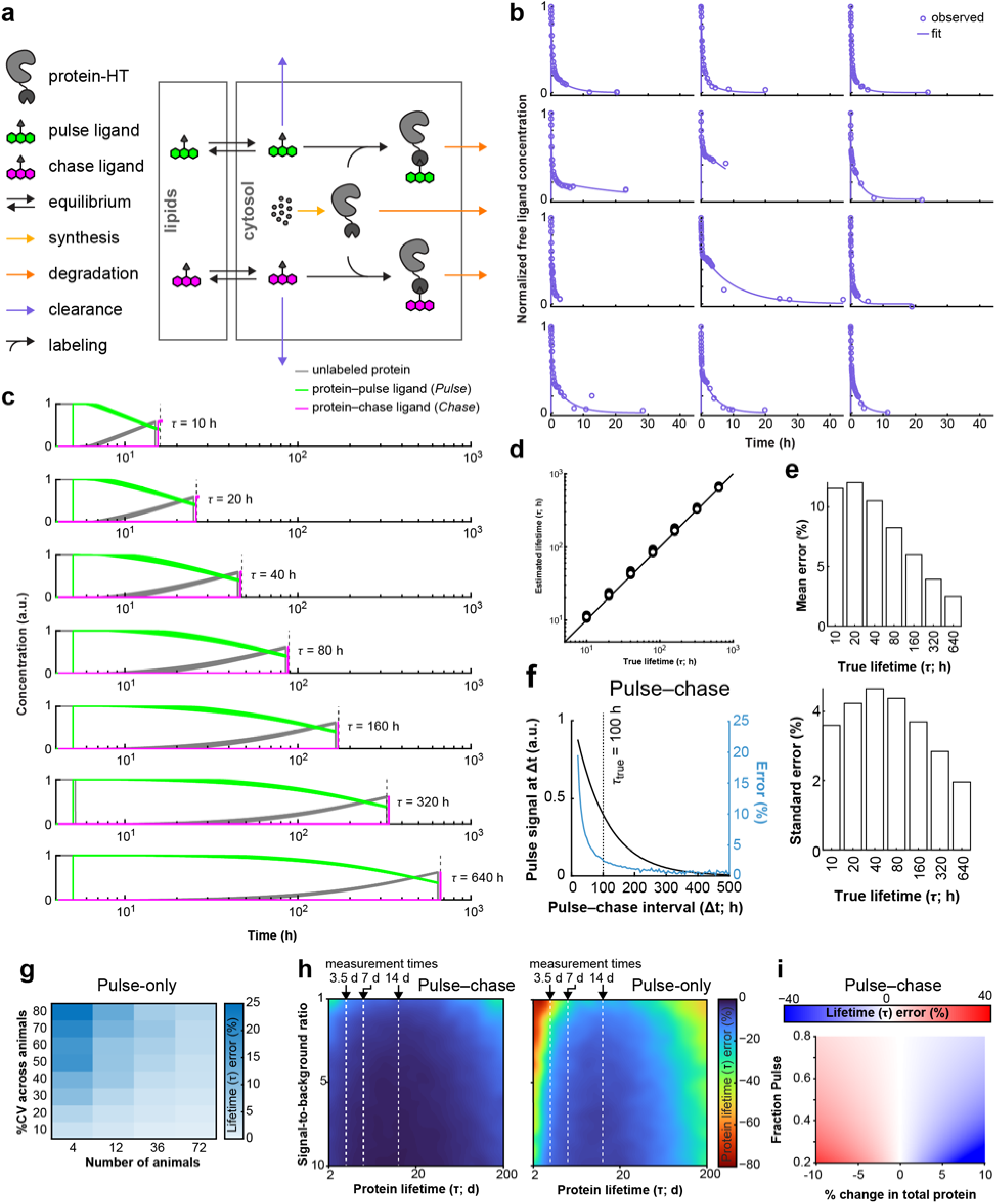
Modeling pulse-chase experiments *in vivo,* constrained by dye clearance measurements. **a**, Model compartments, molecules, and reactions for modeling protein turnover measurements. A pulse or chase dye is added to the cytosol compartment and subsequently: (1) cleared; (2) can move to a lipid compartment (representing the slow dye clearance component); or (3) irreversibly binds to the protein-HT. Here, the protein synthesis rate is equal to its degradation rate, with dye-bound proteins degrading at the same rate. This maintains total protein amount constant regardless of dye binding. See **Supplementary Text** for more information. **b**, Fitting of the dye clearance data (**Extended Data Figure 1**) to generate model variants. Twelve experiments (data in blue circles) and the fitted response (blue lines) are shown. The parameters fit were dye clearance rate, cytosol-to-lipid rate constant, and volume of the lipid compartment. The cytosol compartment was fixed to having a volume of 1 ml. **c-e,** Example model simulations with a selection of 7 HT-protein mean lifetimes (c: 10-640 h; Legend as in Fig. 1a). The correlation between estimated and true protein lifetime is excellent (d). The mean error (e, top) decreases with increasing lifetime, whereas the standard error between model variants (simulating variability in dye clearance) peaks around the mean dye clearance rate (e, bottom). **f**, Simulation of error in the lifetime estimate as a function of the pulse-chase interval (ΔT). The error (blue line) decreases faster than the *Pulse* concentration (black line). This shows that the error with DELTA is dominated by dye clearance, with overall low error rates. **g,** Error in lifetime estimation using pulse-only measurements, given ideal dye injection delivery. Here, variability of the measurement across animals (CV y-axis) could be countered only by averaging across animals (number of animals x-axis). Error rates (color-coded) are higher than in DELTA, where the chase dye and the *Fraction Pulse* calculation normalizes for inter-animal variability under ideal dye injection conditions. **h**, Modeling to compare errors due to shot noise with DELTA (left panel) *vs.* pulse-only labeling (right panel). In both cases three measurement times (white dashed lines) were used to estimate a known decay (x-axis: 2-200 days). Increasing the signal to background ratio (y-axis) and adding a chase dye (as in DELTA; left panel) reduced shot-noise induced errors. **i**, Modeling the effects of a change in total protein on the error in lifetime estimation (% error in colormap) as a function of % change in protein levels (x-axis) and the measured fraction pulse (y-axis). Error increases with longer intervals (smaller fraction pulse) and larger changes in protein.

**Extended Data Figure 3.**
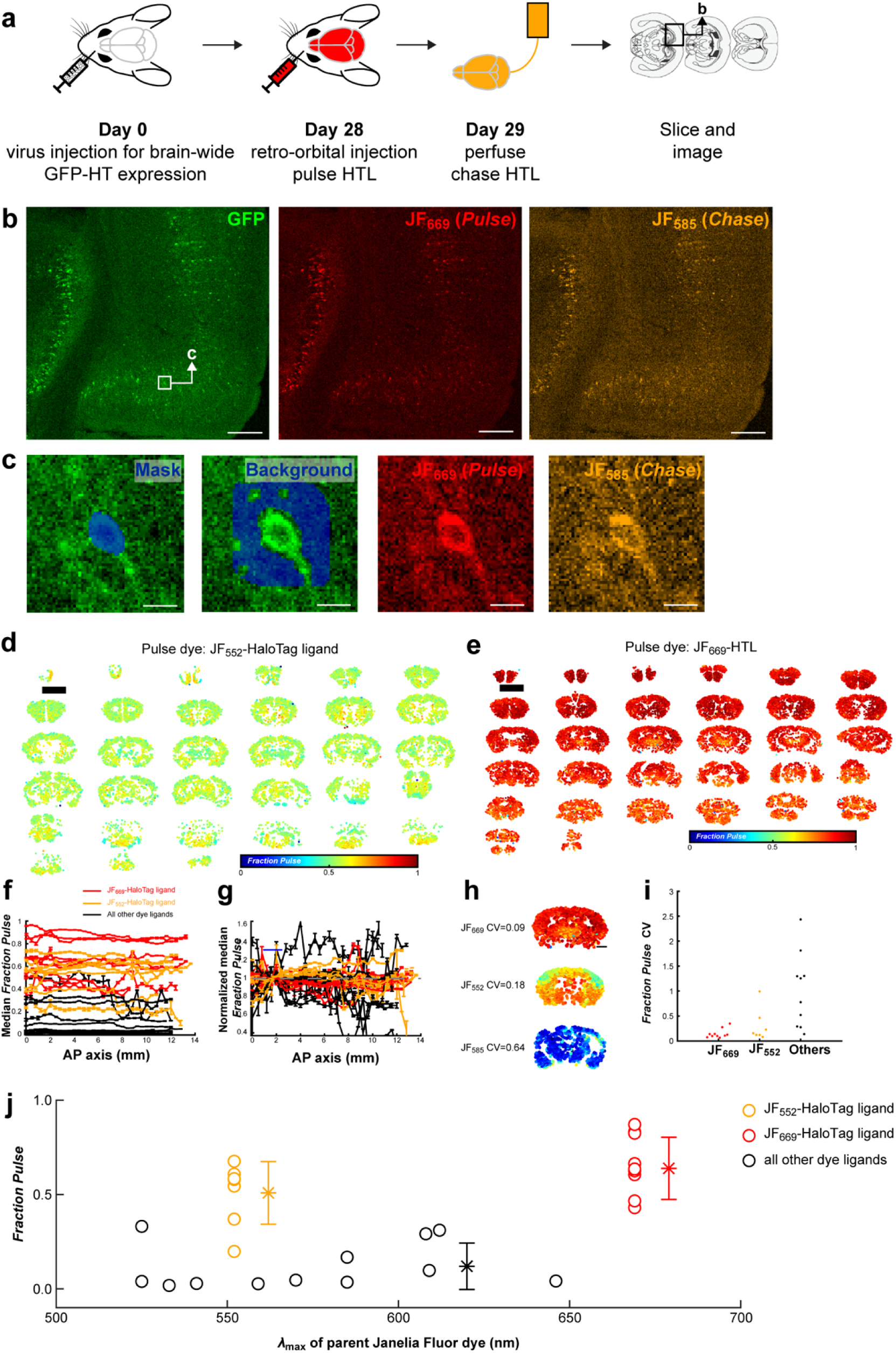
JF_552_ and JF_669_ are bioavailable in the brain. **a,** Experimental procedures for dye screening *in vivo*. **b,** Example coronal sections imaged 4 weeks after infection with an AAV expressing GFP-HT. GFP fluorescence (left panel), fluorescence of JF_669_-HTL injected *in vivo* (*Pulse*; middle panel), and fluorescence of JF_585_-HTL fluorescence applied during perfusion (*Chase*; right panel). Scale bars, 200 µm. **c,** Example cell (square in b, left panel) overlaid with the mask used to extract signal (first panel), a local background (second panel) and the images of the *in vivo* injected dye (third panel) and perfused dye (fourth panel). *Fraction Pulse* is 0.65 in this example. Scale bars are 10 µm. **d,** Coronal slices of GFP-HT injected mouse brain after JF_552_-HTL dye injections. Each cell is colored by *Fraction Pulse*. The scale bar is 3 mm (black bar in the top left). Notice the uniform labeling across the mouse brain. **e,** same as (d) for JF_669_-HTL. **f,** Median *Fraction Pulse* as a function of AP position (of coronal slices). JF_669_-HTL is in red, JF_552_-HTL is in orange, all other HTL dyes are in black. Error bars are standard errors. **g,** After normalization (mean of the median *Fraction Pulse* under the blue line is set to 1), there is no significant deviation from 1 suggesting uniformity of dye distribution in this axis (One-way ANOVA, *F*(19) = 1.46, *p* = 0.0913**). h,** For each coronal section of each animal, a coefficient of variation (CV) was calculated. Three examples of coronal sections are shown. **i,** CV of *Fraction Pulse* for JF_669_-HTL injections (Red, CV = 0.15±0.03, *n* = 10), JF_552_-HTL (Orange, CV = 0.31±0.12, *n* = 7) and the other dyes (Black, CV = 2.2±0.81, *n* = 14). **j,** Mean *Fraction Pulse* for each animal (*n* = 31) injected as a function of the dye excitation wavelength. Here, a higher *Fraction Pulse* signifies better brain bioavailability. JF_669_-HTL (red ; *n* = 10, *Fraction Pulse*: 0.64 ± 0.16) and JF_552_-HTL (yellow; *n* = 7; *Fraction Pulse:* 0.51 ± 0.17) were significantly better than the other dyes (black; *n* = 14, *Fraction Pulse*: 0.12 ± 0.12; 1-way ANOVA *F*(2) = 35.96, *p* = 3.2e-08; post-hoc JF_669_ vs. others *p* = 3.8e-8, JF_552_ vs. others *p* = 2.7e-5).

**Extended Data Figure 4.**
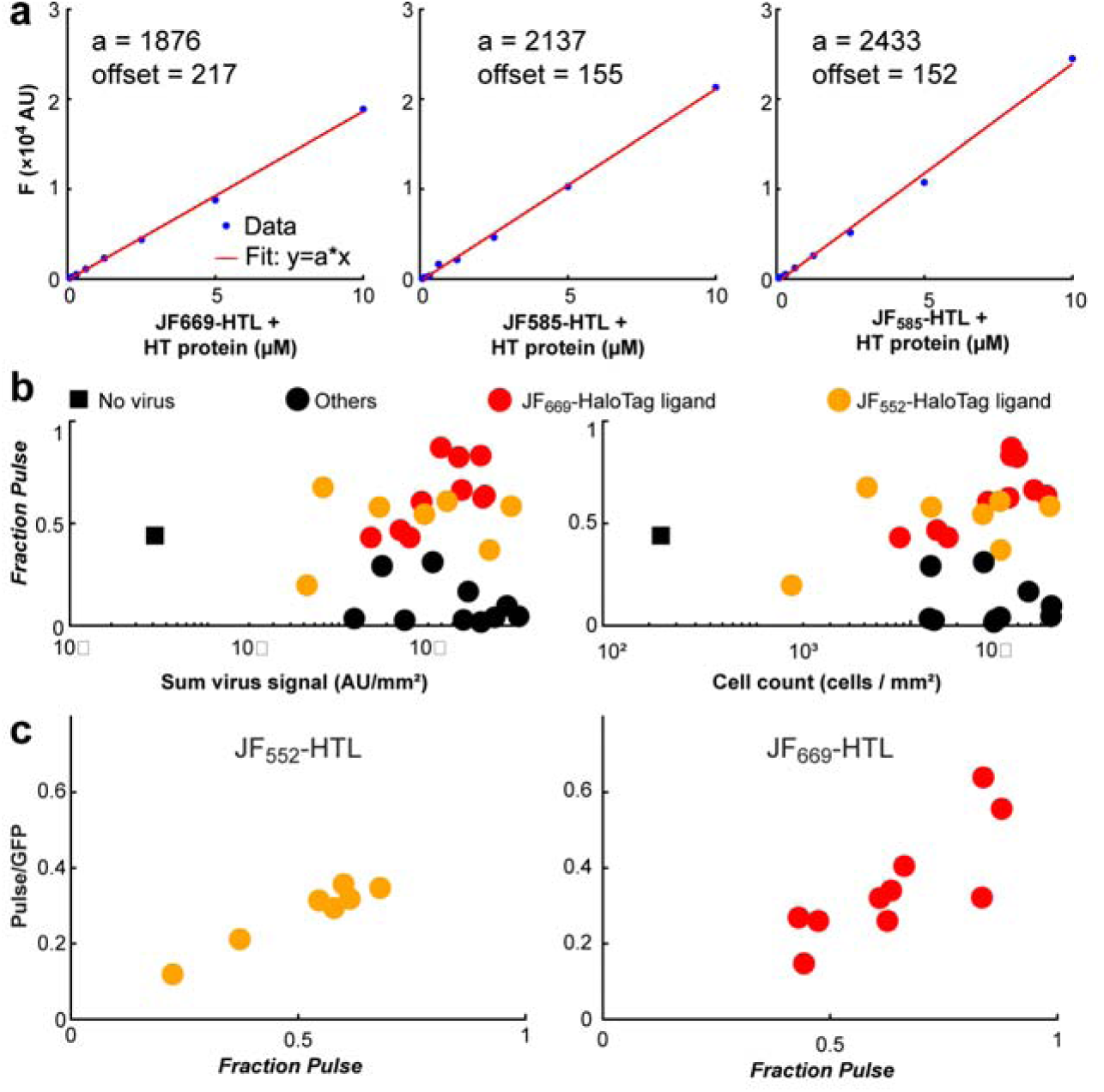
Calibration and validation of GFP-HT based JF dye screening *in vivo.* **a,** Purified HT was added at saturation to a HTL JF dye (left: JF_669_-HTL, middle: JF_585_-HTL, right: JF_552_-HTL) in an 8-well coverslip with 20 μL/well at the following concentrations (μM): 10, 5.0, 2.5, 1.25, 0.625, 0.3125, 0.15625, 0. All 8 wells were imaged under the same conditions as the fixed tissue slides (far red channel for JF_669_ and red channel for JF_552_ and JF_585_). The offset at zero was subtracted and is reported in each panel. A linear slope (red line) was fitted to the data (blue circles) without an intercept term (JF_669_-HTL: *r*^2^ = 0.998; JF_585_-HTL: *r*^2^ = 0.9982; JF_552_-HTL *r*^2^ = 0.9963). This calibration covers 20 out of the 30 animals used and a naive calibration was used for the rest of the dyes (150 offset, 2000 slope). **b,** Left, mean *Fraction Pulse* is not correlated with the sum of the virus signal (GFP channel) per animal (Magenta is JF_669_-HTL, red is JF_552_-HTL, black is other dyes; *r*^2^ = 0.02, *p* = 0.45, *n* = 28). An animal that was not injected with a virus (black square) has at least an order of magnitude less summed fluorescence. Right, mean *Fraction Pulse* is not correlated with the number of cells detected per animal (*r*^2^ = 0.005, *p* = 0.717, *n* = 28; normalized by the area of tissue imaged). An animal that was not injected with a virus (black square) has at least an order of magnitude less detected cells**. c,** As the *Fraction Pulse* increases, so is the ratio between the *Pulse* and GFP, indicating that variability in the *Chase* cannot explain the increases in *Fraction Pulse* (Left: JF_552_-HTL: *n* = 7, *r*^2^ = 0.945, *F*(5) = 85.5, *p* = 0.00025; Right: JF_669_-HTL: *n* = 10, *r*^2^ = 0.65, *F*(8) = 14.7, *p* = 0.00495). **X-axis**: *Fraction Pulse* [*Pulse* / (*Chase* + *Pulse*)]. This compares the pulse injected *in vivo* to the total protein calculated by using the two dyes with their calibration. **<u>Y axis</u>**: *Pulse to GFP ratio.* This compares the pulse injected *in vivo* to the total protein as estimated by the GFP fluorescence. The fact that these 2 ways correlate, means that we are obtaining reliable estimates for the dye calibrations (needed to calculate *Pulse* + *Chase*).

**Extended Data Figure 5.**
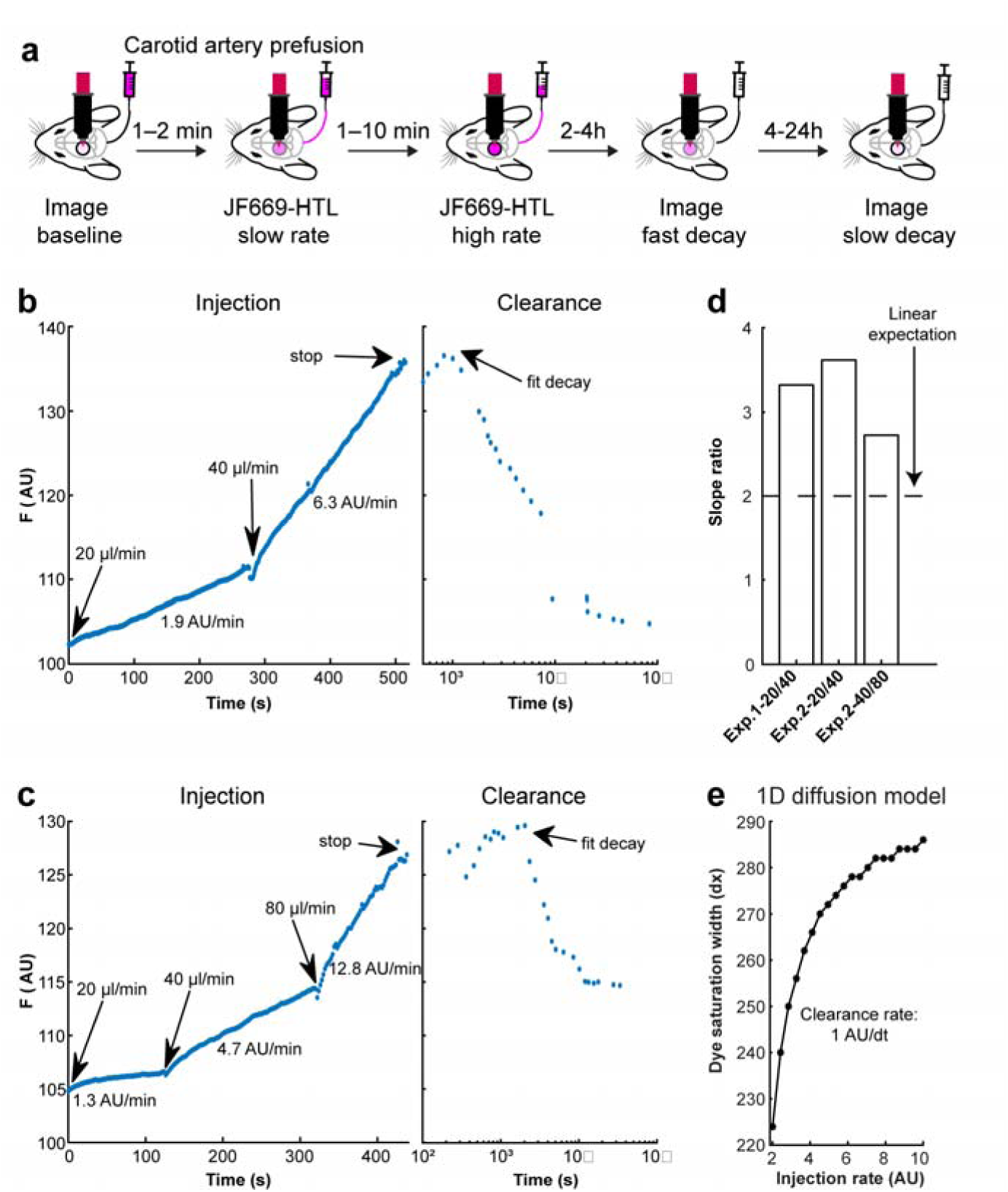
Injection pharmacokinetics of JF dyes. **a,** Design of experiment to measure dye injection pharmacokinetics. Continuous imaging was performed during carotid artery perfusion of JF_669_-HTL at different rates. **b,** One animal, using infusion rates of 20 µl/min and 40 µl/min. **c,** Another animal, using 20, 40, and 80 µl/min. **d,** Slope ratios for all transitions in infusion rates (two animals, three transitions). If clearance was proportional to the amount of dye injected, a slope ratio of 2 is expected. The slope ratio of > 2 implies sublinear clearance or saturation of clearance mechanisms. **e,** 1D diffusion model shows an increase in the area of saturation (y-axis) with increasing dye injection rate (x-axis). See Supplementary Text for further details.

**Extended Data Figure 6.**
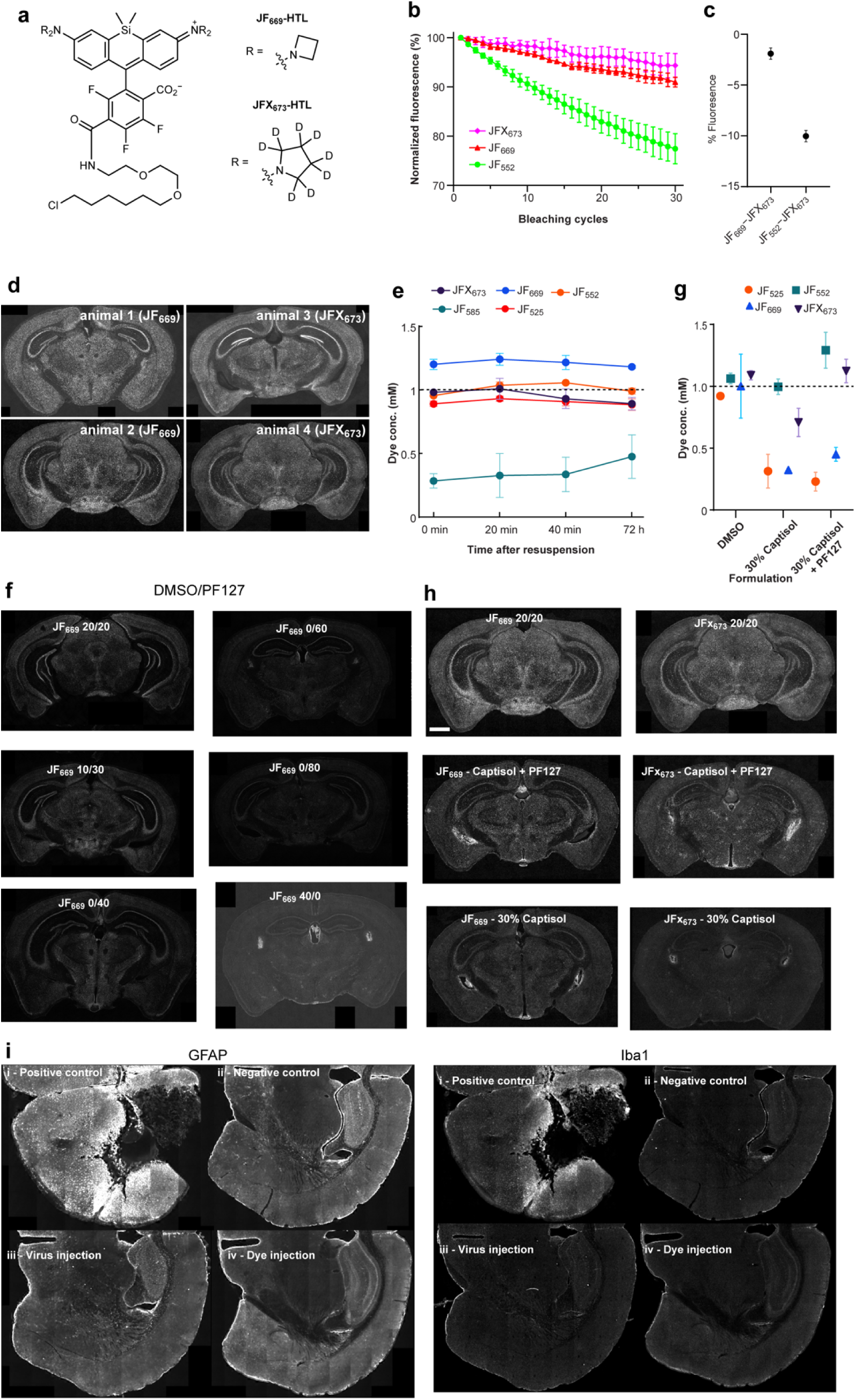
Validation of dye formulation and JFX_673_-HTL, a bioavailable red dye with improved photostability. **a,** Comparison of JF_669_-HTL and JFX_673_-HTL. Note the addition of deuterium and additional carbon in the R position of JFX_673_. **b,** Bleaching curves of JF dyes showing normalized fluorescence over 30 bleaching cycles. **c,** JFX_673_-HTL was significantly more photostable than the other dyes tested. **d,** Example sections of MeCP2-HT mice do not show significant differences in brain availability of JF_669_-HTL versus JFX_673_-HTL. All panels show the pulse dye. **e**, Solubility of JF-HTL dyes using DMSO (20 μl), Pluronic F127 (20 μl), and PBS (60 μl) formulation over time after resuspension. No significant decrease in solubility was observed and all *in vivo* dyes used reached the intended solubility of 1 mM (2-way ANOVA [Dye x Time] p < 0.05; Time: *F*(3,20) = 0.66, *p* = 0.5835; Dye: *F*(4,20) = 149.21, p < 0.0001; Interaction: *F*(12,20) = 1, *p* = 0.4794). **f,** Example coronal slices from MeCP2-HaloTag animals injected with different ratios of DMSO and Pluronic F127. The original 1:1 ratio was the best, as shown by the brighter pulse dye staining. **g,** Different formulation than (e) to test the replacement of DMSO (left) with Captisol (middle) or a combination of Captisol and Pluronic F127 (right). We saw mixed effects on solubility, while some dyes (JFX_673_-HTL) retained solubility, and some did not (JF_669_-HTL; 2-way ANOVA, interaction: *F*(6) = 13.31, p < 0.0001; Dye: *F*(3) = 67.35., *p* < 0.0001; Formulation: *F*(2) = 46.91, *p.*< 0.0001). **h,** Example coronal slices from MeCP2-HaloTag animals injected with solutions from (f). Both dyes that were less soluble with Captisol alone or Captisol with Pluronic F127 (JF_669_ left) or those that were as soluble (JFx_673_ right) were less bioavailable without DMSO. **i,** Immunohistochemistry against GFAP (left) and Iba1(right) for 4 conditions, i-Positive control: A cortical lesioned animal. ii-Negative control: Naive animal without any manipulation. iii-Virus injection: An animal was injected with the GFP-HaloTag virus and perfused 3 weeks after injection. iv-Dye injection: An animal was injected with JF_669_-HTL and perfused 24 h after injection. Both dye injection and virus injection do not increase GFAP or Iba1 levels.

**Extended Data Figure 7.**
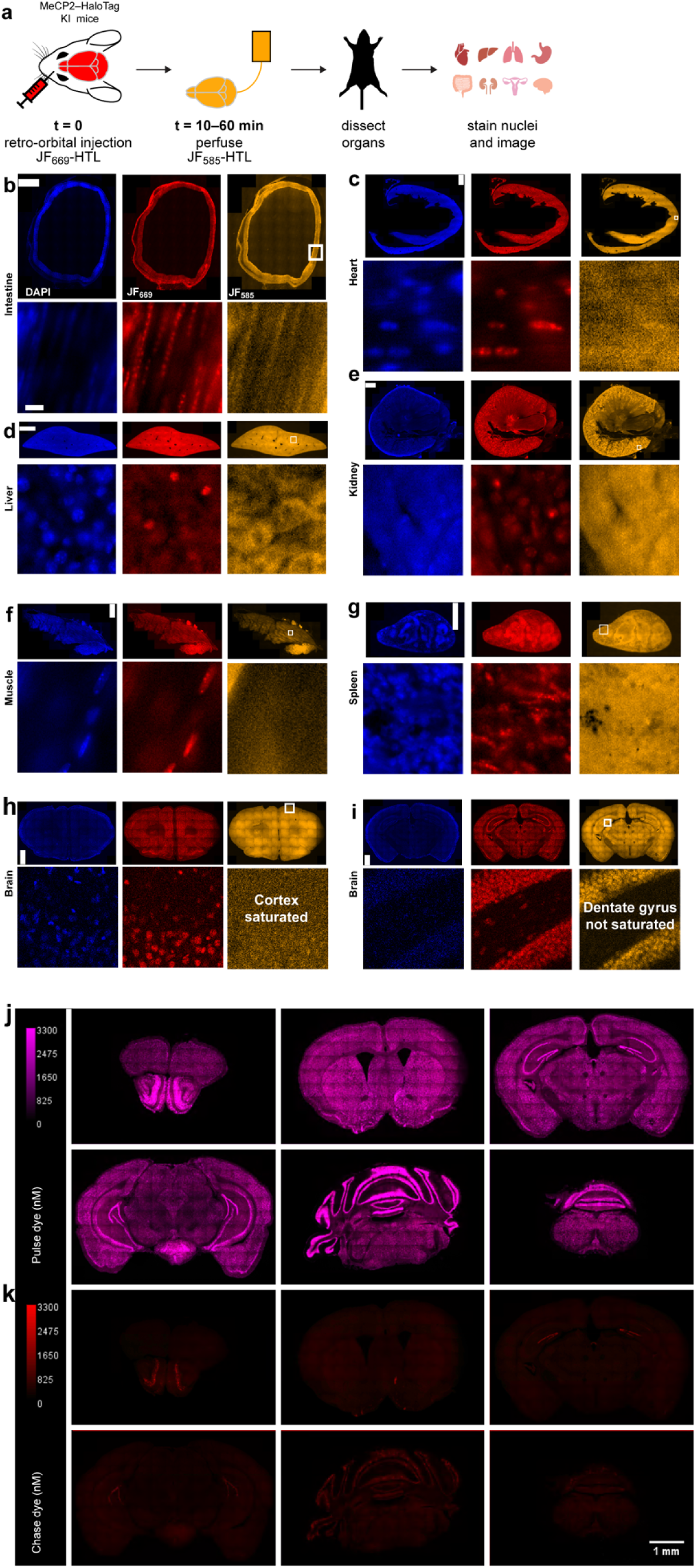
Systemic injection of ligand dye saturates the abundant protein MeCP2-HT in most organs. **a,** Schematic of the experimental procedures to test the saturation of MeCP2-HT. **b-i,** Organs harvested and imaged for pulse dye saturation (red), as evident by the lack of chase dye (orange) in the nuclei (blue) where MeCP2-HT is expressed. Top scale bar is 1 mm, bottom scales bars are 50 μm. **j-k,** Six coronal sections of an MeCP2-HT animal injected *in vivo* with JF_669_-HTL (j) immediately followed by perfusion with JF_552_-HTL (k) to check for saturation. All panels were converted to nM dye concentration and scaled to the same concentration. Only very dense cell body regions, highly enriched in MeCP2 (*e.g.*, i), show any chase dye staining, indicating the inability to saturate *in vivo*.

**Extended Data Figure 8.**
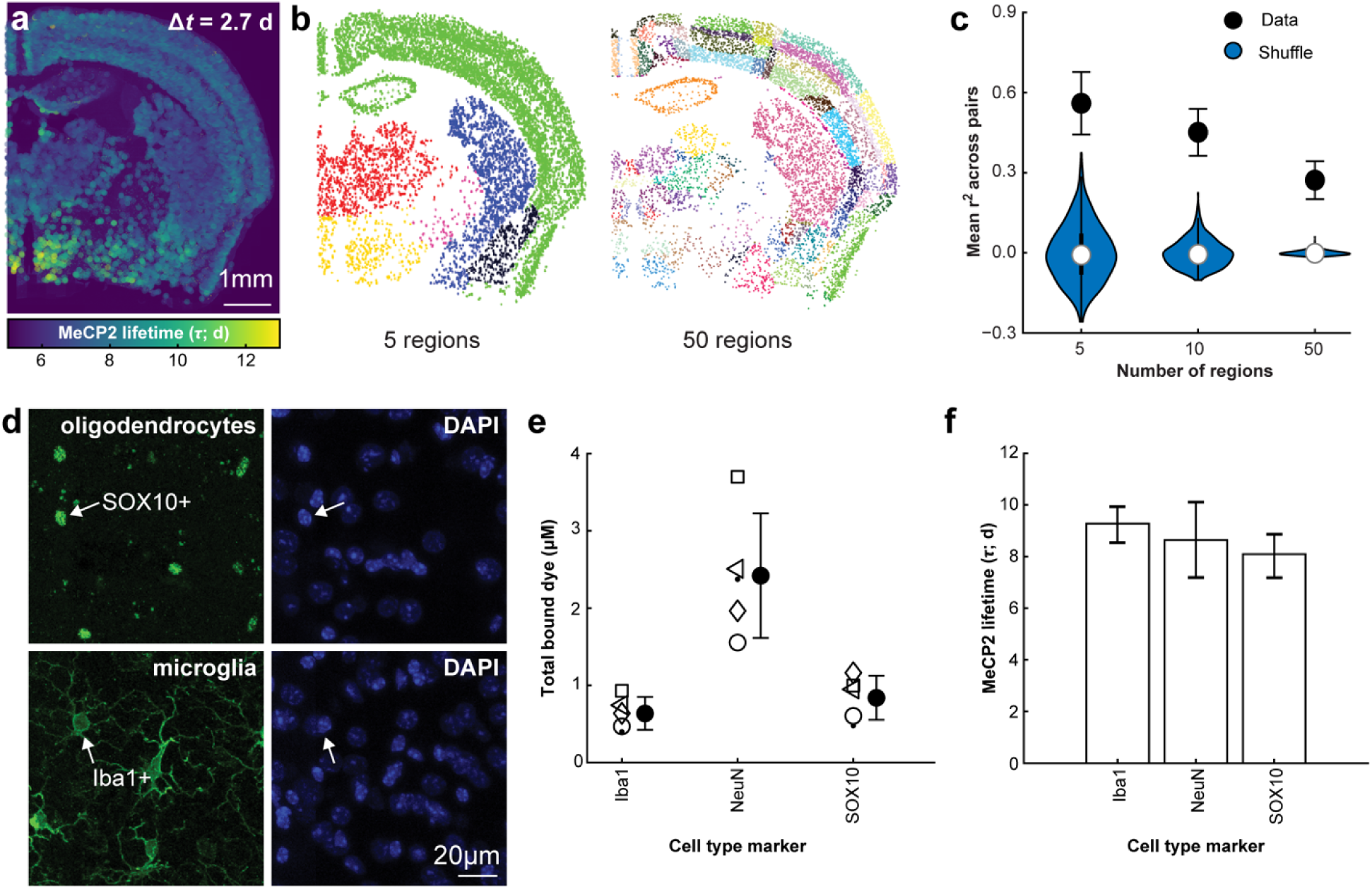
The measured lifetime of MeCP2-HT is consistent across individual mice and cell-types. **a,** Example coronal section of the lifetime of MeCP2-HT after segmentation of NeuN positive nuclei. **b,** Example assignment of neuronal nuclei to brain regions based on the Allen CCF v3 at different levels (left 5 regions, right 50 regions). **c,** Protein lifetime measurements were consistent between mice as assessed by examining correlations between all possible pairs of animals (*n* = 5 animals, 10 pairs). For three levels of the CCF (5, 10, and 50 regions) the correlation between regions was higher than for a shuffled control. **d,** Two additional sections were stained in each of the five animals for oligodendrocytes (SOX10, top) and microglia (Iba1, bottom). **e,** MeCP2 was more abundant in the nuclei of neurons (Iba1: 0.63 ± 0.21, NeuN: 2.42 ± 0.8 SOX10: 0.84 ± 0.28 µM total dye signal; ANOVA *F*(2) = 27.22, *p* = 2e-4). **f,** Lifetimes were similar for all cell types, as confidence intervals overlapped between all cell types (mean and [CI] of Iba1: 9.2 [8.5-9.9], NeuN: 8.6 [7.1-10.0], SOX10: 8.1 [7.2-8.9] days of MeCP2 lifetime).

**Extended Data Figure 9.**
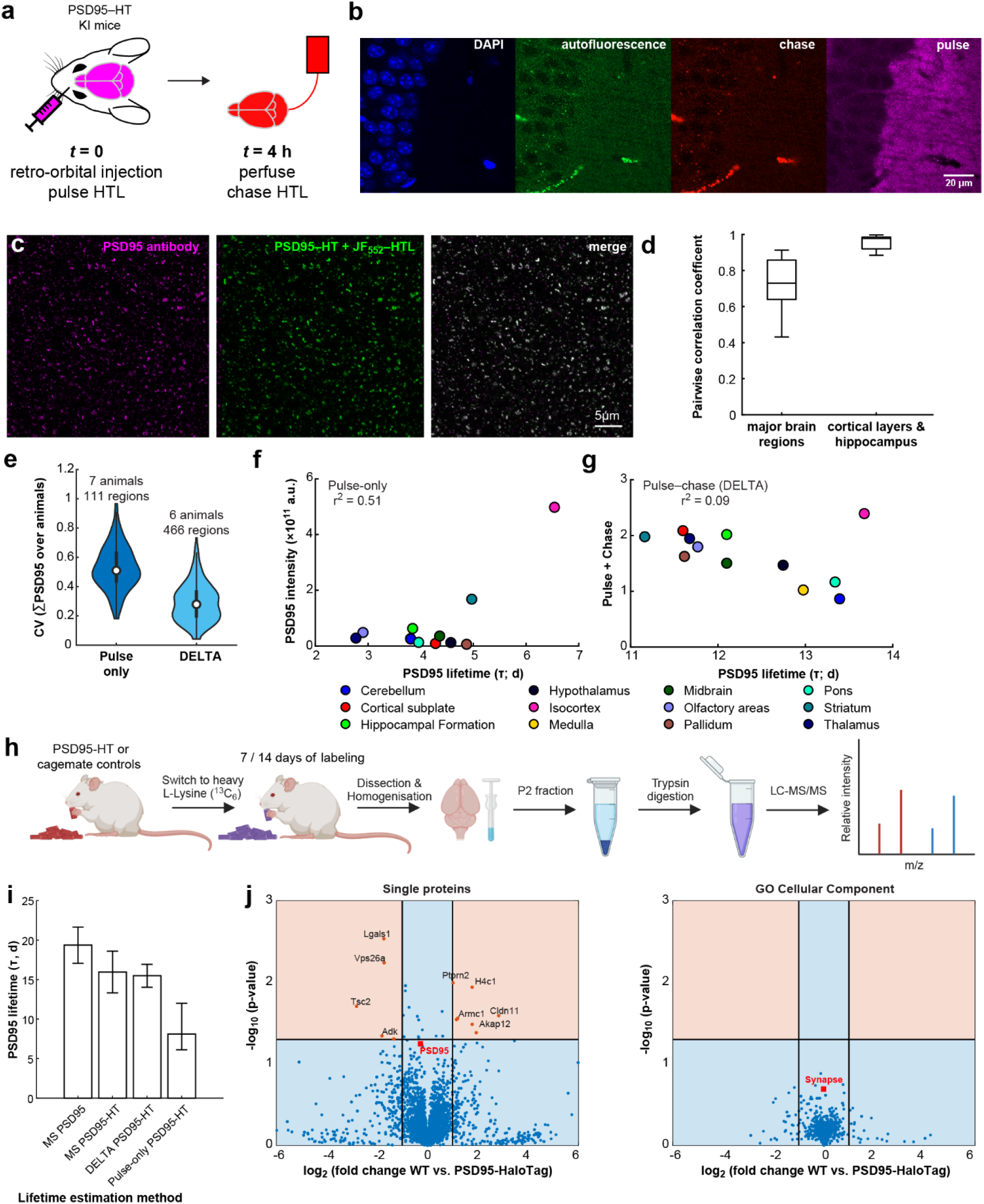
Validation of DELTA in the PSD95-HaloTag mouse. **a,** Experimental procedures for testing saturation in a PSD95-HT mouse. **b** Example section showing the basal dendrite section of the CA1 region of the hippocampus (DAPI shows the nuclei in the leftmost panel). The green channel (autofluorescence; 2^nd^ panel from left) accounts for all the signal in the red channel (*Chase* – JF_552_-HTL; 3^rd^ panel), while the far-red channel shows the expected pattern of synapses (*Pulse* – JFX_673_-HTL; 4^th^ panel). **c**, Validation of HTL signal (JF_552_-HTL; middle panel) in the PSD95-HT knock-in animal with a knockout validated antibody (left panel) showing high colocalization (right panel). **d**, Box plot of all pairs of animals (6 animals, *n* = 15 pairs) for correlation coefficients for the lifetime of PSD95 across 12 large brain regions (left panel; As in Fig. 2g**,i**) or cortical layers and HC subfields (right; As in Fig. 2f**,j**). **e**, Violin plot of coefficient of variation (CV) across animals for the total PSD95 measured for each brain region. Pulse only is publicly available data from Bulovaite et al. (2022) where we used the standard deviation divided by the mean of integrated fluorescence at day 0 (*n* = 7 animals; *n* = 111 brain regions). For DELTA we used the standard deviation divided by the mean of the total *Pulse* and *Chase* values (*n* = 6 animals; *n* = 466 brain regions). Measurement variability of total PSD95 across animals was lower in DELTA then in Bulovaite et al. (Wilcoxon rank sum test; *z* = 12.6; *p* = 1.3e-36). **f-g**, Correlation between estimated expression of PSD95 (y axis) with its lifetime measurement (x axis) from data of Bulovaite et al. (f; Pulse only) or from DELTA (g) for twelve brain regions (colored circles). r^2^ values from linear regression are shown for each panel. See Supplementary Text for details. **h**, Workflow for mass spectrometry (MS) measurements of protein turnover. **i**, Comparisons of PSD95 lifetime with or without HaloTag fusion by different methods. There is an agreement between DELTA and MS based measurements of lifetime for PSD95-HT. The non-overlapping confidence intervals of the pulse-only approach signifies a statistically significant difference. **j**, Volcano plots comparing MS-based proteome-wide lifetime estimates between PSD95-HaloTag knock-in mice and WT cage mates. Left, per protein analysis showing most proteins don’t significantly change in lifetime (y-axis: -log_10_ *p* value; horizontal line at *p* = 0.05; x-axis: log_2_ fold change; vertical lines at two-fold increase or decrease). PSD95 is highlighted with a red square, significant single proteins are labeled with their gene names. Right, same as left but averaging proteins based on their GO Cellular Component annotations. Synapse Cellular Component is highlighted with a red square.

**Extended Data Figure 10.**
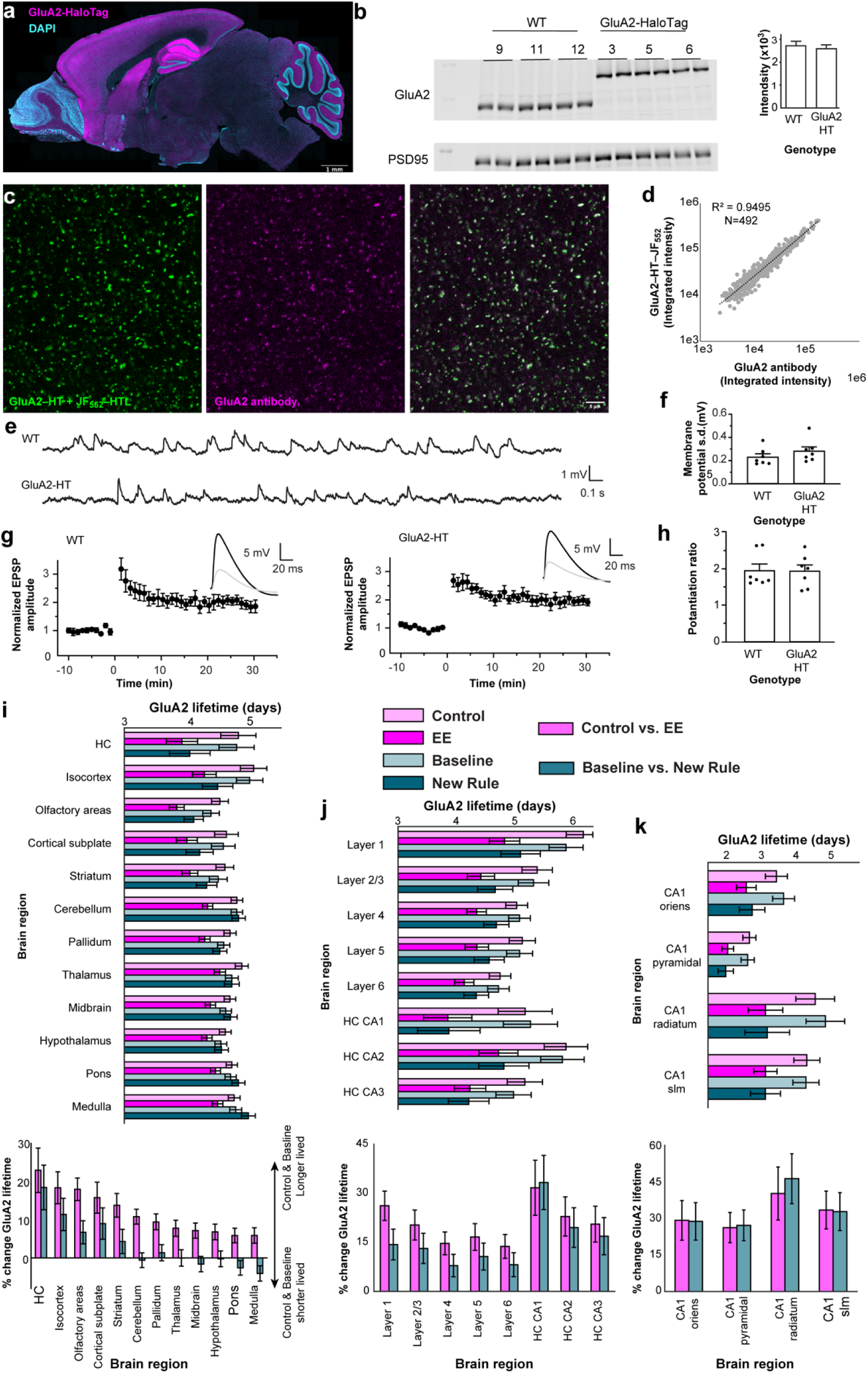
GluA2-HaloTag validation for use in DELTA brain-wide. **a**, Sagittal section for a GluA2-HT knock-in mouse perfused with JFx_673_-HTL dye showing the distribution of GluA2 (magenta). Nuclei were stained with DAPI (white). **b**, Left, Western blots stained for GluA2 and PSD95 as normalization. 12 lanes were used: 3 WT animals (Lanes marked: 9,11,12) and 3 homozygous GluA2-HT knock-ins (Lanes marked: 3,5,6) both done in duplicates. Right, quantification of GluA2 signal normalized to PSD95 averaged across animals and replicates for WT (27,186±2004) and GluA2-HT knock-ins (25,963±1685). **c**, Validation of HTL signal (JF_552_-HTL; LEFT panel) in the GluA2-HT knock-in animal with an antibody (middle panel) showing high colocalization (right panel). **d**, Segmentation of 492 synapses from (c) quantifying the integrated intensity in the HTL-dye channel (y axis, log scale) vs. the antibody signal (x-axis, log scale) showing very high correlation (r^2^ = 0.9495) **e**, Example spontaneous activity traces of WT (top) and knock-in (bottom) from acute HC slices. **f**, Quantification of spontaneous activity as the standard deviation of the traces show no significant difference (WT: 0.23±0.03 mV, knock-in: 0.28±0.04 mean ± SE, *n* = 7 for both, *p* = 0.29 2-sided T-Test). **g**, Example LTP induction in WT (top) and knock-in (bottom). **h**, Quantification of the potentiation ratio shows normal LTP of the knock-in compared to WT (WT: 1.96 ± 0.17 knock-in: 1.98±0.18 mean ± SE, *n* = 7 for both, *p* = 0.94 2-sided t-test). **i-k**, Average lifetimes (top) of the 4 behavioral conditions (Control, EE, Baseline, New Rule) and 2 comparisons of lifetime (bottom; Control vs. EE and Baseline vs. New Rule) for the 12 large brain regions (i) neocortical layers and HC subfields (j) and for CA1 lamina (k). All statistics in i-k come from a Mixed Effects model in which group assignment is a fixed effect and animal IDs are random effects.

## Supplementary text

### Simulations of pulse-chase experiments

We used simulations to gain an understanding of the conditions that would enable DELTA to provide precise measurements of protein turnover. We investigated multiple sources of error, including dye pharmacokinetics, amount of dye injected, variability in dye clearance, pulse-chase interval (ΔT), and compared DELTA to a pulse-only paradigm^28^. While the dye amount and ΔT are experimentally controlled, a crucial unknown is the pharmacokinetics of the HTL dyes. For example, if the free HTL dye is not cleared quickly, newly synthesized HT-protein could be labeled by the pulse dye instead of the chase dye, thus biasing the measurement to longer lifetimes (gray region in **Fig. 1a**).

We first measured the rate of dye clearance in the brains of wild-type (WT) mice. HTL JF dyes were injected into the retro-orbital sinus (**Extended Data Figure 1a; Methods**)^72^ and imaging was subsequently performed with a wide-field fluorescence microscope^71^ through a cranial glass window^90,91^, for up to 24 hours after injection (**Extended Data Fig. 1b** for example imaging session). Dye concentration decayed with a dominant fast component (14±2 min), and a smaller amplitude slow component (197±37 min; **Extended Data Fig. 1c-d** and **Supplementary Table 1**; fast/slow amplitude ratio: 1.7±0.3; *n* = 12; corresponding to a geometric average dye lifetime of 82 min). We incorporated dye clearance kinetics into our model (**Extended Data Fig. 2a**). Dye in a cytosolic compartment could either bind to the free protein-HT, directly clear out (corresponding to the fast component), or partition into a second compartment (lipids; corresponding to the slow component). We measured dye clearance in 12 mice. Fitting model parameters to each experiment (**Extended Data Fig. 2b**) produced 12 model variants that differ in the three parameters that impact dye clearance kinetics (lipids compartment volume, fast clearance rate constant, and cytosol to lipids equilibrium constant).

Because each HT-protein exists at different concentrations *in vivo*, it is difficult to inject the precise amount of HT ligand to reach saturation without leaving free ligand. Thus, should we aim to under-saturate or provide dye excess? To address this question, we varied the ratio of dye to protein-HT target and estimated the error in estimating protein turnover for a broad range of average protein lifetimes (**Fig. 1b**). The dependence of the error on protein lifetime has three regimes. First, under-saturated labeling (**Fig. 1b** - dye/protein ratio < 1) results in large errors, regardless of protein lifetime. Second, dye saturation (**Fig. 1b** - dye/protein ratio > 1) results in large errors for short protein lifetimes, mirroring the under-saturation regime. Third, dye saturation results in small errors for long protein lifetimes (>40h). The main factor contributing to this effect in the dye saturating regime is the time of dye clearance relative to the measurement time. As the dye clears within an hour (**Extended Data Fig. 1**), the relative contribution to the error increases for shorter lived proteins, which entail shorter pulse-chase intervals. Specifically, in the saturating regime (>1.2 dye/protein ratio), dye clearance kinetics would bias the lifetime estimate towards larger values, as slower dye clearance results in labeling of proteins synthesized after the pulse injection (gray shaded region in **Fig. 1a**). This would increase *Pulse* values and decrease *Chase* values, thus inflating the lifetime estimation. In a mild saturation regime (1-1.2 dye/protein ratio) during dye injection, new protein is made for shorter-lived proteins, thus more dye is consumed during the dye clearing window, which reduces the error introduced by dye excess. These competing factors create a nonmonotonic relationship between the estimation error and the protein lifetime (**Extended Data Fig. 2e**). Thus, two of the necessary conditions for reliably measuring the protein lifetime using DELTA are saturation of the target protein with pulse dye injection while avoiding very short-lived proteins. This is not very limiting as most proteins in the brain have lifetimes much longer than the one hour dye-clearance time^3,12^.

We next measured the effect of model variants (combination of the lipids compartment volume, fast dye clearance rate, and cytosol-to-lipids equilibrium constant) on the estimation of protein lifetime across a wide range of simulated lifetimes (**Extended Data Fig. 2c)**. These variants capture the uncontrolled variable corresponding to the variance in dye clearance that would lead to variance (not bias) in our measurements. Using a pulse-chase interval equal to the average protein lifetime (ΔT == r) and a pulse dye amount in the mildly saturating regime (1.2 dye/protein ratio), we compared the estimated lifetime with the true one and found a good correspondence (**Extended Data Fig. 2d**; *r*^2^ = 0.9996 *n* = 84, 12 variants × 6 lifetimes). We identified a nonmonotonic relationship between bias and protein lifetime (Mean of error as defined above; **Extended Data Fig. 2e** – left panel), as expected from the single-model variant simulation. Of note, while bias (a shift of the mean) would lead to an overestimation of the lifetime, it does not affect our ability to make comparisons between brain regions in the same animal or between animals. Dye injection variability (Standard deviation of Error; **Extended Data Fig. 2e** – right) would affect our ability to compare the lifetime and has a nonmonotonic shape as well. However, in all cases these were small (less than 5%).

### Comparing DELTA with other protein turnover methods

We first compared DELTA with MS. MS results vary between individual studies depending on the cellular fraction selected and other experimental conditions (*e.g.*, *in vitro vs. in vivo*; for a review see Alvarez-Castelao & Schuman^92^), exhibiting considerable variability for any single protein, making comparisons across animals difficult. In our own experiments we observed that the HT fusion reduces PSD stability by approximately 2 days (**Extended Data Fig. 9i**), explaining some of the differences between DELTA and previous MS measurements. Additional differences might be related to the challenges in accounting for the recycling of isotope-labeled amino acids in MS experiments. A key advantage of DELTA over MS is the lack of recycling. Due to the enzymatic activity of HT on the HTL, a HT-protein that is degraded would release a ligand that can’t rebind to a different unlabeled protein. Furthermore, as the chase HTL dye is delivered during perfusion, no degradation of the *Chase* is expected.

Next, we looked at other fluorescent imaging methods, which allow for labeling with different fluorophores at specific time points by fusing a tag with a protein of interest. This approach offers better spatial resolution, but remains difficult to deploy in the brain due to issues with epitope tags and labeled antibodies^93,94^ or the self-labeling SNAP-tag^95^ system and cognate ligands that do not cross the blood-brain barrier^96,97^. Lastly, other expression methods such as in *utero* electroporation to overexpress HT^55^ or local injection of viruses to knock-in HT^98^ does not allow sufficient spatial sampling to compare brain-wide turnover rates across mice.

We also compared DELTA to single-dye (pulse-only) methods^28,96^. First, we estimated the effects of the pulse-chase interval (Δt) in DELTA on the lifetime estimation error. While the protein decayed exponentially (**Extended Data Fig. 2f** – black line), error decreased faster as function of Δt (**Extended Data Fig. 2f** – blue line). This decay of error again supported the idea that the main bias in this measurement is the relative time of dye clearance from the measurement time. Here, only one animal is modeled, so no averaging is used to calculate turnover rates. In contrast, pulse-only methods require multiple animals to determine the initial vs. later protein content of any given brain region. We modeled pulse-only experiments by assuming the same lifetime (100 h), a perfect measurement of the *Pulse*, and that the only variability would be across animals. We assumed that different animals would have on average 1 AU HT-protein and varied the coefficient of variation (CV 10-80%; **Extended Data Fig. 2g** – y axis). The range was selected conservatively according to the values seen in mass spectroscopy (MS) for biological replicates^99,100^. Variability could be reduced by averaging across multiple animals. We calculated the error in estimating the lifetime as a function of the number of animals used (**Extended Data Fig. 2g – x axis;** 4-72). These were distributed over four time points (0, 10, 30, 100h) in which a single exponential decay was fitted after averaging across animals (average number of animals per time point of 1, 3, 6, 12 animals). The estimation error increased with increasing protein expression variability and decreased with averaging over animals, as expected. However, for most operating regimes of DELTA, there would be no error under these conditions, as the *Fraction Pulse* calculation normalizes for the total protein amount.

We also investigated another source of error, photon shot noise. In pulse-only methods, lower expressing regions would reach the noise floor faster than higher expressing regions as *Pulse* signal declines with time. We simulated a pulse-only vs. pulse-chase experiments where signal to background ratio (SBR) was used as a proxy for expression strength (**Extended Data Fig. 2h)**. In each simulation we assumed perfect dye delivery and no variability in protein expression. A baseline of 100 photons was used as a noise floor (to avoid counting errors). A protein (τ=2-200h; **Extended Data Fig. 2h** – x axis) was simulated using a signal of 100-1000 photons at t=0 representing SBR of 1-10 (**Extended Data Fig. 2h** – y axis) and measurements were taken at t=3.5, 7, 14 h. Mean errors of protein lifetimes were computed using bootstrapping and assuming Poisson noise. As expected, higher SBR reduced errors regardless of protein lifetime or methodology. However, DELTA was better in all simulated conditions (**Extended Data Fig. 2h)**. This has the prediction that low expression of a protein might have a negative effect on the ability to measure its turnover. Indeed, in publicly available data from Bulovaite et al.^28^ there is a correlation between protein expression at day 14 and measured protein lifetime (**Extended Data Fig. 9f**). In contrast, we don’t see such a relationship with the sum of the *Pulse* and *Chase* in DELTA (**Extended Data Fig. 9g**).

Finally, DELTA offers an improved ability to estimate changes in protein lifetime, even when there is a change in the total protein amount across experimental conditions. In DELTA, the *Pulse* + *Chase* accurately measures the total protein amount, which helps avoid errors in turnover estimation that would otherwise arise from unaccounted changes in total protein (**Extended Data Fig. 9i**).

### Screening for bioavailable JF dyes

We screened HTL JF dyes that would be able to saturate abundant proteins in the brains of mice by measuring bioavailability in the brain (**Extended Data Fig. 3**). Our target was green fluorescent protein tagged with HT (GFP-HT). AAV-PHP.eB expressing GFP-HT was introduced by retro-orbital injection^76^, which led to sparse and brain-wide expression. Four weeks after viral transduction, HTL dyes (**Supplementary Table 2**) were injected retro-orbitally. After 12-18 hours, we perfused the brain with a spectrally orthogonal dye. Given that GFP has a lifetime of several days^101^, *Fraction Pulse* is a measure of the dye’s ability to saturate the protein-HT target in the live animal.

The brain was sectioned coronally and imaged with a confocal slide scanner. Example images from an injection show the target protein in green (**Extended Data Fig. 3b**, left panel), the *in vivo* delivered dye in red (JF_669_-HTL; **Extended Data Fig. 3b**, middle panel) and perfusion dye (JF_585_-HTL; **Extended Data Fig. 3b**, right panel) in orange. This procedure enabled segmentation and analysis of single cells (**Extended Data Fig. 3c**). The fluorescence values of individual cells were background corrected and converted to dye concentration using calibration curves (**Extended Data Fig. 4a**). These calibrations are needed to determine the total protein amount (*Pulse* + *Chase*) correctly as the illumination and detection sensitivities of the dyes vary.

We identified two dyes that saturated GFP-HT in the brain: JF_669_-HTL and JF_552_-HTL. These dyes were significantly better than the other dyes tested (**Extended Data Fig. 3j**). Variability in GFP-HT expression did not account for the variability in these dyes, as the number of detected cells or virus signals did not correlate with *Fraction Pulse* (**Extended Data Fig 4b**). We also expect that a higher *Fraction Pulse* value would correlate with a higher *Pulse*/GFP ratio. Here, a large difference would implicate issues with the perfusion and complete saturation of the GFP-HT target with the *Chase*. However, *Fraction Pulse* did correlate with the *Pulse*/GFP ratio, as expected if the perfusion dye saturates the remaining GFP-HT proteins. This was the case both for JF_669_-HTL and JF_552_-HTL (**Extended Data Fig. 4c**).

We next quantified the amount of variability across brain regions in the delivery of the dye under these under-saturation conditions. We examined individual cells in all coronal sections imaged (examples in **Extended Data Fig. 3d,e**). These injections produced low variability along the AP axis (**Extended Data Fig. 3f**), as indicated by normalizing the average *Fraction Pulse* in the front of the brain and the absence of any trend toward the back of the brain (**Extended Data Fig. 3g**). Looking at individual coronal sections (**Extended Data Fig. 3h**), for our bioavailable dyes we see a low CV for each slice (**Extended Data Fig. 3i**). This indicates low variability in dye delivery across the brain for JF_669_-HTL and JF_552_-HTL which would facilitate identification of variability in protein lifetime across brain regions (**Extended Data Fig. 3j**).

To assess successful saturation in a challenging case (MeCP2 is a very abundant protein, making it challenging to saturate in vivo), we injected JF_669_-HTL retro-orbitally (*Pulse*) to the MeCP2-HT animal. This was followed by perfusion with JF_585_-HTL (*Chase*) after 1 h (**Extended Data Fig. 7a**). The lack of *Chase* indicated saturation with the pulse dye ligand in almost all tissues, including most regions of the brain (**Extended Data Fig. 7b–h**). Some brain regions containing dense cell body layers were not saturated, however, including the hippocampal CA1 region (**Extended Data Fig. 7i**). Nuclei located farther from the vasculature exhibited lower labeling intensities, suggesting this is due to ligand depletion in regions with densely clustered nuclei expressing large amounts^41^ of a HT fusion protein. Nevertheless, we achieved MeCP2–HT saturation in most brain regions (**Extended Data Fig. 7j,k**).

To assess the consistency of our method with MeCP2-HT, we looked at pairwise correlations across animals for all commonly imaged brain regions. The average mean correlation across pairs was high compared to shuffled controls for three levels of region annotation representing different spatial scales based on the Allen Brain Reference common coordinate framework (CCFv3; **Extended Data Fig. 8a–c**)^49^.

Finally, we did not see a spatial correlation between the brain vasculature^102^ and turnover measurements using the PSD95-HT animal. For example, in both the hippocampus and neocortex, most blood vessels are oriented dorsoventrally while the pattern of turnover is mediolateral in the hippocampus (CA1 *vs.* CA3) and dorsoventral in neocortex (Layer 1 to Layer 6; **Fig. 2e**). This approach provides a good test case for biases that could arise from inhomogeneous delivery of ligands via the vasculature, but we found that PSD95 signal is homogenous at this resolution. As with MeCP2, we looked at pairwise correlations across animals for all commonly imaged brain regions. PSD95-HT turnover was highly consistent across mice as the average mean correlation across pairs was high (**Extended Data Fig. 9d**).

### Formulation and injection of bioavailable JF dyes

Janelia Flour (JF) HaloTag ligand (HTL) dyes are a key reagent for DELTA. We characterized these dyes in terms of infusion kinetics, formulation, solubility, and the brain’s reaction in terms of inflammation. We first wanted to understand whether the injection rate can affect the dye’s ability to saturate proteins in the brain. However, due to the retro-orbital injection procedure, we were unable to capture the dye dynamics during injection as the mouse headbar physically interfered with the retroorbital injection while it was mounted for imaging. Additionally, it is difficult to precisely control the rate of delivery in retroorbital injections, as they are performed manually. To perform these types of experiments we needed a precise i.v. dye delivery method that would enable simultaneous brain imaging. We choose to use carotid artery perfusion to deliver JF_669_-HTL at different rates during continuous imaging through a cranial window (**Extended Data Fig. 5a**). In these experiments, if dye clearance is linearly proportional to the dye injection rate, then doubling the rate of dye delivery should exactly double the rate of dye increase in the brain. This would tell us that the dynamics of dye injection would not dramatically change the total amount of dye delivered to the brain. However, if the slope of dye accumulation in the brain increases more than twofold, it would indicate a saturation of the dye clearance mechanism. This would favor faster dye injections leading to higher peak concentrations and total dye delivery to the brain. In the first mouse (**Extended Data Fig. 5b**) we started perfusion at 20 µl/min and later increased to 40 µl/min **(Supplementary Movie 3**). Increasing the delivery rate resulted in a more than twofold increase in the slope of dye accumulation in the brain (1.9 to 6.3 AU/min). We repeated this experiment with another animal (**Extended Data Fig. 5c)** and obtained similar results **(Extended Data Fig. 5d**). We validated our intuition about the effects of a constant clearing rate using simulations. Using the same amount of dye (20 AU) and a constant clearance rate (1 AU/dt), we simulated different injection rates (2-10 AU/dt). We observed saturation further away from the injection center for faster injection rates (**Extended Data Fig. 5e**). These results favor an injection method that delivers the dye faster given a constrained amount of dye.

To understand our dye constraints, we checked the dye solubility. We validated that our dyes are soluble at our desired concentration and stable over time (1 mM; **Extended Data Fig. 6e**). We used the published formulation for injection^71^, which consists of 20 µl DMSO, 20 µl Pluronic F127, and 60 µl PBS. If we are near the solubility limit, the amount of dye in solution could decrease over time. This was not the case for all dyes injected *in vivo* (**Extended Data Fig. 6e**). The only dye that did not reach our solubility goal was JF_585_-HTL, so it was not used *in vivo*. It was used during perfusion at a much lower concentration (1 µM) well below the solubility limit (∼250 µM).

We then wanted to know if the ratio of DMSO to Pluronic F127 (1:1 20 µl each) was optimal for dye availability. We tested both increasing and reducing the ratio, but the original ratio was the best (**Extended Data Fig. 6f**). As we were injecting 40-80 µl (∼2-4 g/kg) of DMSO, which is lower than the LD50 (>10 g/kg)^103^ but not a small amount, we tried to replace DMSO with Captisol as cosolvent and evaluated both the solubility and the in vivo injections. We saw mixed effects of dyes and formulations by looking at solubility after a 3-day incubation with DMSO, Captisol and a combination of Pluronic F127 and Captisol (**Extended Data Fig. 6g**). We tested both JF_669_ and JFX_673_ using retro-orbital injections with different formulations. As expected, from the solubility data, JF_669_ in other formulations was less bioavailable (**Extended Data Fig. 6h** – left column). However, although JFX_673_ was soluble in all formulations, it was still less bioavailable in the brain using Captisol (**Extended Data Fig. 6h** – Right column).

Finally, we validated that there were no signs of brain inflammation by staining for GFAP and Iba1 markers that we validated with a cortically lesioned animal (**Extended Data Fig. 6i** - panel i) and a naïve animal (**Extended Data Fig. 6i -** panel ii). Both retro-orbital virus injections (**Extended Data Fig. 6i -** panel iii) and our chosen dye injection formulation (**Extended Data Fig. 6i -** panel iv) did not affect these measures. This indicates that there is no major breach of the brain blood barrier, as it is known to induce this type of inflammation^104^.

**Supplementary Table 1.**
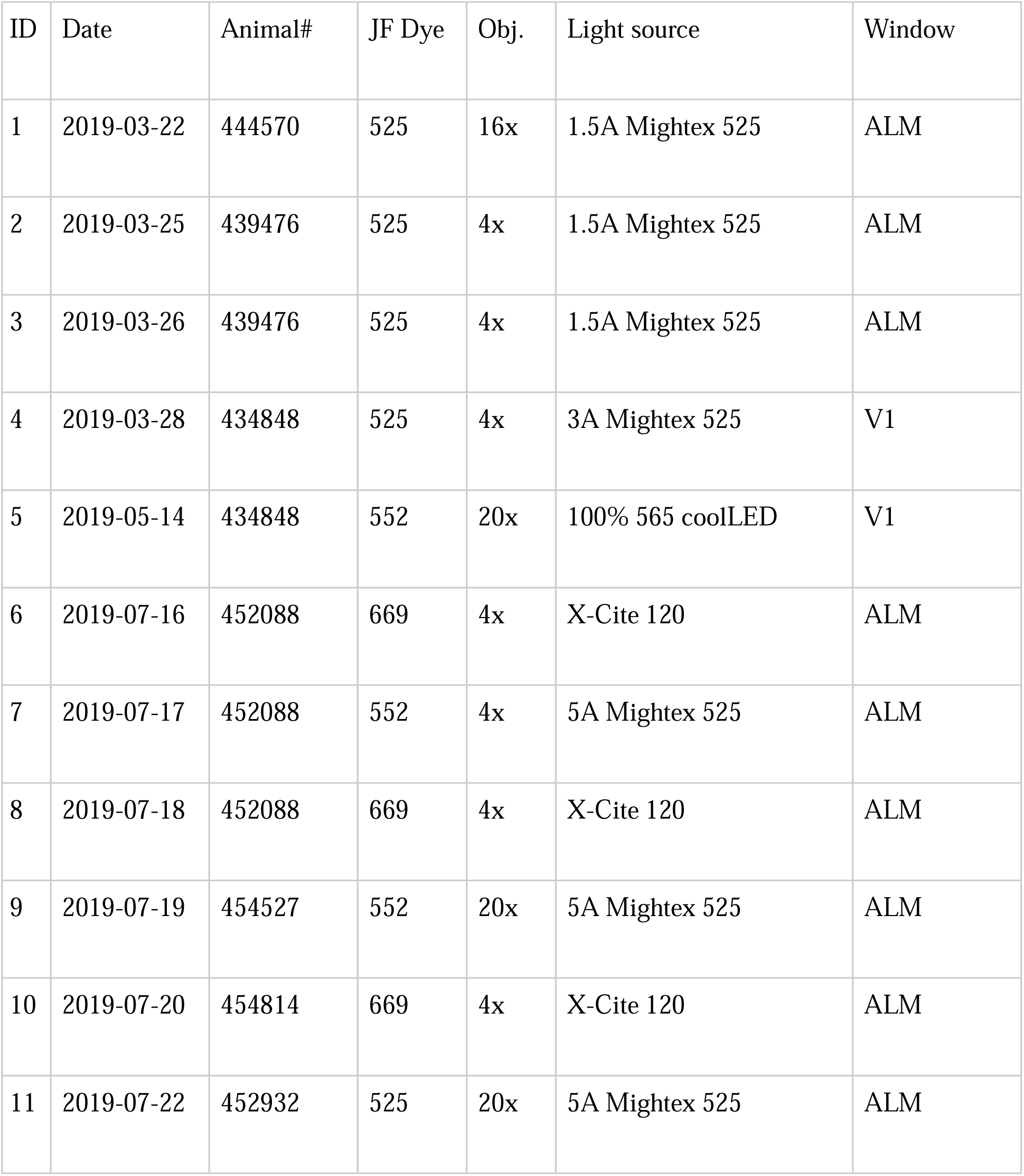
Animals and filters used for dye clearance experiments.

**Supplementary Table 2:** Animals and dyes used for ex vivo screening.

| Batch | Animal number | DOB | Virus injection date | HT-Expression | Dye injection | In-vivo dye | DMSO/PF127/Saline/Injection (μl) | Chase dye | Mount | Fraction *in vivo* |
| --- | --- | --- | --- | --- | --- | --- | --- | --- | --- | --- |
| 2 | 439359 | 8/14/18 | 11/26/18 | GFP | 1/3/19 | JF669 | 20/20/200/200 | JF585 | 5th | 0.47 |
| 2 | 439360 | 8/14/18 | 11/26/18 | GFP | 1/3/19 | JF669 | 20/20/400/200 | JF585 | 5th | 0.43 |
| 3 | 444182 | 10/16/18 | 1/3/19 | GFP | 2/5/19 | None | virus only | None | 1 | hist. |
| 3 | 444185 | 10/16/18 | 1/3/19 | GFP | 2/5/19 | JF585 | 30/10/200/200 | JF669 | 5th | 0.04 |
| 3 | 444186 | 10/16/18 | 1/3/19 | GFP | 2/5/19 | JF585 | 10/30/200/200 | JF669 | 5th | 0.17 |
| 5 | 447231 | 12/4/18 | 4/5/19 | GFP | 5/1/19 | JF669 | 20/20/200/200 | JF585 | 4th | 0.64 |
| 5 | 447232 | 12/4/18 | 4/5/19 | GFP | 5/1/19 | JF669 2x | 20/20/200/200 | JF585 | 4th | 0.66 |
| 5 | 447233 | 12/4/18 | 4/5/19 | GFP | 5/1/19 | JF669 2x | 20/20/200/200 | JF585 | 4th | 0.61 |
| 5 | 447234 | 12/4/18 | 4/5/19 | GFP | 5/1/19 | JF669 | 10/30/200/200 | JF585 | 4th | 0.82 |
| 5 | 447235 | 12/4/18 | 4/5/19 | GFP | 5/1/19 | JF669 | 10/30/200/200 | JF585 | 4th | 0.43 |
| 6 | 445766 | 12/5/18 | 4/12/19 | GFP | 5/8/19 | JF552 2x | 20/20/200/200 | JF669 | 4th | 0.54 |
| 6 | 445767 | 12/5/18 | 4/12/19 | GFP | 5/8/19 | JF552 2x | 20/20/200/200 | JF669 | 4th | 0.58 |
| 6 | 445768 | 12/5/18 | 4/12/19 | GFP | 5/8/19 | JF552 | 20/20/200/200 | JF669 | 4th | 0.68 |
| 6 | 447239 | 12/4/18 | 4/12/19 | None | 5/8/19 | JF552 | 20/20/200/200 | JF669 | 4th | hist. |
| 6 | 447240 | 12/4/18 | 4/12/19 | GFP | 5/8/19 | JF552 2x | 20/20/200/200 | JF669 | 4th | 0.61 |
| 7 | 452036 | 2/19/19 | 4/19/19 | GFP | 5/18/19 | JF541 | 20/20/200/200 | JF669 | 24th | 0.03 |
| 7 | 452037 | 2/19/19 | 4/19/19 | GFP | 5/18/19 | JF559 | 20/20/200/200 | JF669 | 24th | 0.03 |
| 7 | 452038 | 2/19/19 | 4/19/19 | GFP | 5/18/19 | JF533 | 20/20/200/200 | JF669 | 24th | 0.02 |
| 7 | 452039 | 2/19/19 | 4/19/19 | GFP | 5/18/19 | JF552 | 20/20/200/200 | JF669 | 4th | 0.37 |
| 8 | 452031 | 2/19/19 | 4/26/19 | GFP | 5/22/19 | JF669 | 20/20/200/200 | JF585 | 4th | 0.87 |
| 8 | 452032 | 2/19/19 | 4/26/19 | GFP | 5/22/19 | JFX612 | 20/20/200/200 | JF585 | 4th | 0.31 |
| 8 | 452033 | 2/19/19 | 4/26/19 | GFP | 5/22/19 | 552 | 20/20/200/200 | JF669 | 4th | 0.58 |
| 8 | 452034 | 2/19/19 | 4/26/19 | GFP | 5/22/19 | 608 | 20/20/200/200 | JF552 | 4th | 0.29 |
| 8 | 452035 | 2/19/19 | 4/26/19 | GFP | 5/22/19 | Jfx608 | 20/20/200/200 | JF552 | 4th | 0.1 |
| 10 | 451116 | 3/11/19 | 7/15/19 | mKate2 | 8/12/19 | 525 | 20/20/200/200 | JF669 | 4th | 0.33 |
| 10 | 451119 | 3/11/19 | 7/15/19 | mKate2 | 8/12/19 | 525 | 20/20/200/200 | JF669 | 4th | 0.04 |
| 11 | 458564 | 5/21/19 | 8/8/19 | GFP | 11/11/19 | JF669 | 20/20/200/200 | JF585 | 4th | 0.83 |
| 11 | 458565 | 5/21/19 | 8/8/19 | GFP | 10/24/19 | JF669 | 20/20/200/200 | JF552 | 4th | 0.63 |
| 11 | 458566 | 5/21/19 | 8/8/19 | GFP | 10/24/19 | JF646Bio | 20/20/200/200 | JF552 | 4th | 0.04 |
| 11 | 458567 | 5/21/19 | 8/8/19 | GFP | 11/11/19 | JF552 | 20/20/200/200 | JF669 | 4th | 0.2 |
| 11 | 458568 | 5/21/19 | 8/8/19 | GFP | 10/24/19 | JF570 | 20/20/200/200 | JF669 | 4th | 0.05 |
| 13 | 460147 | 7/13/19 |  | MeCP2-F | 1/15/20 | JF669 | 40/0/60/100 | JF585 | 24th |  |
| 13 | 460149 | 7/13/19 |  | MeCP2-F | 1/15/20 | JF669 | 20/20/60/100 | JF585 | 24th |  |
| 13 | 467293 | 6/1/19 |  | MeCP2-M | 1/15/20 | JF669 | 40/0/60/100 | JF585 | 24th |  |
| 13 | 467294 | 6/1/19 |  | MeCP2-M | 1/15/20 | JF669 | 20/20/60/100 | JF585 | 24th |  |
| 13 | 467295 | 6/1/19 |  | MeCP2-M | 1/15/20 | JF669 | 40/0/60/100 | JF585 | 24th |  |
| 13 | 467296 | 6/1/19 |  | MeCP2-M | 1/15/20 | JF669 | 20/20/60/100 | JF585 | 24th |  |
| 14 | 456996 | 6/1/19 |  | MeCP2-F | 2/10/20 | JF669 | 20/20/60/100 | JF585 | 24th |  |
| 14 | 456997 | 6/1/19 |  | MeCP2-F | 2/10/20 | JF669 | 0/40/60/100 | JF585 | 24th |  |
| 14 | 456998 | 6/1/19 |  | MeCP2-F | 2/10/20 | JF669 | 10/30/60/100 | JF585 | 24th |  |
| 14 | 456999 | 6/1/19 |  | MeCP2-F | 2/10/20 | JF669 | 0/60/40/100 | JF585 | 24th |  |
| 14 | 457000 | 6/1/19 |  | MeCP2-F | 2/10/20 | JF669 | 0/80/20/100 | JF585 | 24th |  |

**Supplementary table 3:** MS results comparing WT to PSD95-HT mice protein turnover in the cortex.

**Supplementary Movie 1 – Imaging turnover of single synapses.** Imaging single synapses in layer 1 of cortex and in CA3 subfield of the hippocampus using ExM and Airyscan imaging. Related to Figure 2k.

**Supplementary Movie 2 – Intracellular pool of GluA2-HT in cortex.** In green is the extracellular pool labeled by JF_549i_-HTL. In magenta is the intracellular pool labeled with JFX_673_-HTL. In cyan is nuclei stained with DAPI. Related to Figure 4d.

**Supplementary Movie 3 – Carotid artery infusion**. Carotid artery infusion of dye at a slow rate (20 µl/min) followed by at a faster rate (40 µl/min). Related to Extended Data Fig. 5.

